# Base editing strategies to convert CAG to CAA diminish the disease-causing mutation in Huntington's disease

**DOI:** 10.1101/2023.04.28.538700

**Authors:** Doo Eun Choi, Jun Wan Shin, Sophia Zeng, Eun Pyo Hong, Jae-Hyun Jang, Jacob M. Loupe, Vanessa C. Wheeler, Hannah E. Stutzman, Benjamin P. Kleinstiver, Jong-Min Lee

## Abstract

An expanded CAG repeat in the huntingtin gene (*HTT*) causes Huntington’s disease (HD). Since the length of uninterrupted CAG repeat, not polyglutamine, determines the age-at-onset in HD, base editing strategies to convert CAG to CAA are anticipated to delay onset by shortening the uninterrupted CAG repeat. Here, we developed base editing strategies to convert CAG in the repeat to CAA and determined their molecular outcomes and effects on relevant disease phenotypes. Base editing strategies employing combinations of cytosine base editors and gRNAs efficiently converted CAG to CAA at various sites in the CAG repeat without generating significant indels, off-target edits, or transcriptome alterations, demonstrating their feasibility and specificity. Candidate BE strategies converted CAG to CAA on both expanded and non-expanded CAG repeats without altering *HTT* mRNA and protein levels. In addition, somatic CAG repeat expansion, which is the major disease driver in HD, was significantly decreased by a candidate BE strategy treatment in HD knock-in mice carrying canonical CAG repeats. Notably, CAG repeat expansion was abolished entirely in HD knock-in mice carrying CAA-interrupted repeats, supporting the therapeutic potential of CAG-to-CAA conversion base editing strategies in HD and potentially other repeat expansion disorders.

## Introduction

Huntington’s disease (HD; MIM #143100) ^1–3^ is one of many trinucleotide repeat disorders caused by expansions of CAG repeats ^4–8^. Although the underlying causative genes, pathogenic mechanisms, clinical features, and target tissues may be different ^7; 9; 10^, these disorders share a cardinal feature: an inverse relationship between age-at-onset and respective CAG repeat length ^4; 7; 11–19^. To explain this striking genotype-phenotype correlation that is common to many trinucleotide repeat expansion disorders, a universal mechanism in which length-dependent somatic repeat expansion occurs toward a pathological threshold has been proposed ^20^. This mechanism provides a good explanation of the relationship between CAG repeat length and age-at-onset in HD very well as 1) the *HTT* CAG repeat shows increased repeat length mosaicism in the target brain region ^21–23^, 2) somatic instability is repeat length-dependent ^23; 24^, and 3) the levels of repeat instability shows correlations with cell type-specific vulnerability and age-at-onset ^22; 23; 25^. In addition, somatic repeat instability of an expanded *HTT* CAG repeat appears to play a major role in modifying HD since our genome-wide association studies have revealed that the majority of onset modification signals represent instability-related DNA repair genes ^26–29^. Together, these data support the critical importance of CAG repeat length and somatic instability in determining the timing of HD onset.

Recent large-scale genetic analyses of HD subjects have revealed that different DNA repeat sequence polymorphisms have an impact on age-at-onset. Most HD subjects carry an uninterrupted glutamine-encoding CAG repeat followed by a glutamine-encoding CAA-CAG codon doublet (referred to as a canonical repeat; CR) ^24; 27; 30^. However, expanded CAG repeats lacking the CAA interruption (loss of interruption; LI) or carrying two consecutive CAA-CAG (duplicated interruption; DI) ^24; 27; 30^ also exist (S. Figure 1). Surprisingly, the age-at-onset of HD subjects carrying LI or DI alleles is best explained by the length of their uninterrupted CAG repeat, not the encoded polyglutamine length ^24; 27; 30^. These human genetics data indicate that introducing CAA interruption(s) into the *HTT* CAG repeat to reduce the length of the uninterrupted repeat is a potential therapeutic avenue to delay the onset of HD. Importantly, a genome engineering technology called base editing (BE) was recently developed, permitting the C-to-T conversion (cytosine base editors; CBEs) or A-to-G conversion (adenine base editors; ABEs) ^31–34^, where CBEs could in principle be applied to convert CAG codons to CAA to shorten the uninterrupted CAG repeat without altering polyglutamine length or introducing different amino acids. In view of the strong human genetic evidence for the role of the uninterrupted CAG repeat length in determining HD onset, we have conceived BE strategies of converting CAG codons to CAA within the repeat and evaluated their therapeutic potential in HD.

**Figure 1.**
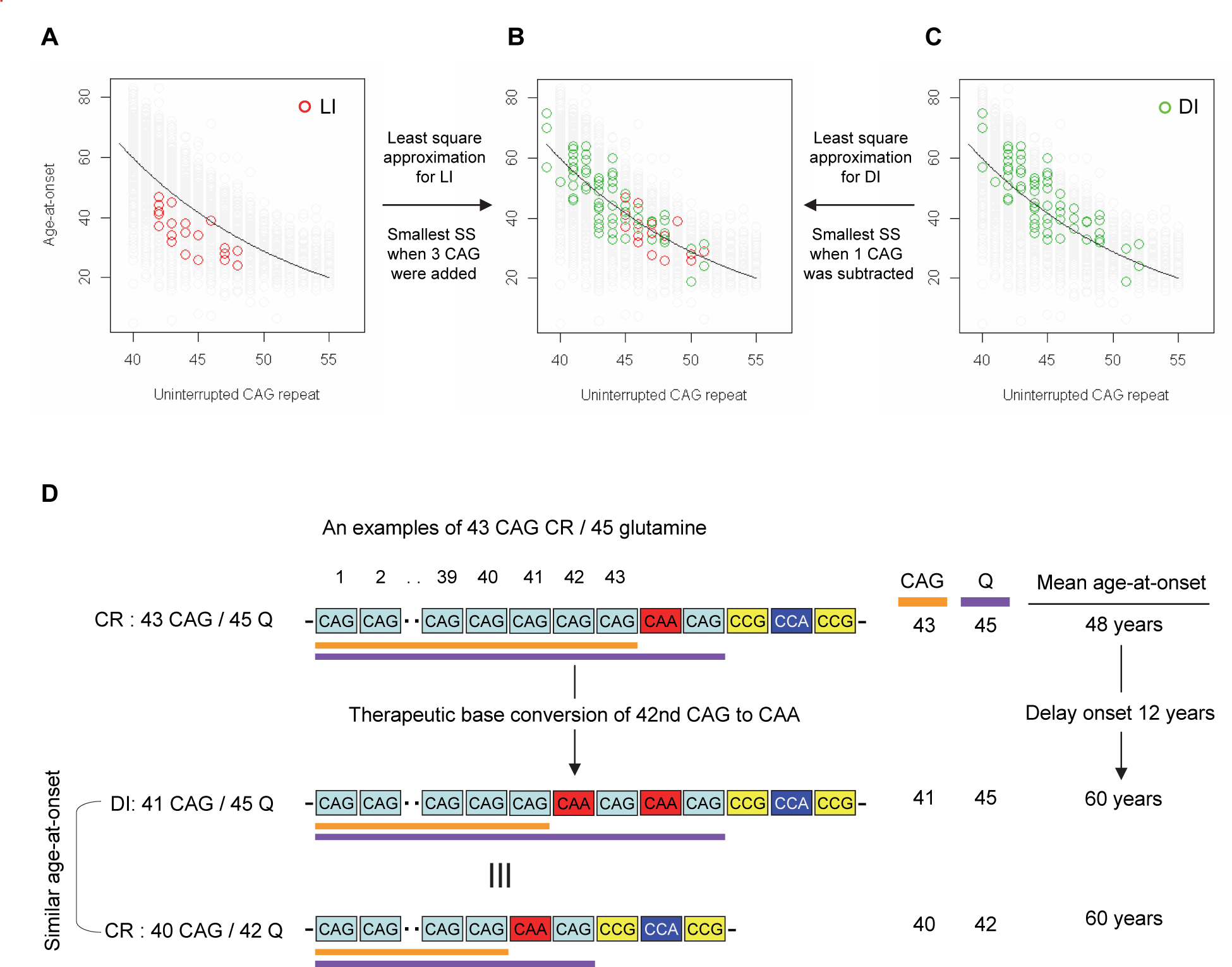
Effects of CAA interruption on HD age-at-onset. (A-C) Least square approximation was performed to estimate the additional effects of LI (red circles in panel b and c) and DI on age-at-onset (green circles in panel c and d). We varied the CAG length of HD participants carrying LI or DI, and subsequently calculated sum of square (SS) to identify the CAG repeat that explained the maximum variance in age-at-onset of these allele carriers. Y-axis and X-axis represent age-at-onset and CAG repeat length, respectively. Grey circles and black trend lines respectively represent HD participants with CR alleles and their onset-CAG relationship. SS means sum of square. (D) To illustrate the magnitude of the impact of a therapeutic base editing strategy of converting a CR allele to a DI allele by changing CAG to CAA, an example of a CR of 43 CAG (45 glutamine) with a mean observed onset of 48 years is displayed. In this example, therapeutic conversion of the 42nd CAG to CAA by BE would produce a DI allele of 41 CAG (45 glutamine). Considering the additional effect of DI alleles in HD patients, a 41 CAG / 45 glutamine DI allele would produce an onset similar to a CR allele of 40 CAG / 42 glutamine, with a mean onset age of 60. Therefore, CAG-to-CAA conversion in HD subjects with 43 CR repeats could delay onset by 12 years.

## Material and Methods

### Study approval

Subject consents and the overall study were approved by the Mass General Brigham IRB and described previously ^27^. Experiments involving mice were approved by the Mass General Brigham Institutional Animal Care and Use Committee.

### Age-at-onset of HD subjects carrying LI or DI

Detailed experimental procedures for sequencing of the *HTT* CAG repeat region and determination of the CAG repeat length are described previously ^27^. We compared age-at-onset of HD subjects carrying LI or DI to that of HD subjects carrying CR. Expected age-at-onset from CAG repeat length of CR was based on the onset-CAG regression model that we reported previously ^19; 26; 27^. For expected age-at-onset based on the polyglutamine length, the same regression model was modified by replacing CAG repeat length with CAG repeat length plus 2 because the glutamine length equals CAG + 2 in CR.

### Least square approximation to estimate the additional effects of LI and DI on age-at-onset

Carriers of LI and DI alleles showed slightly earlier and later onset age, respectively, compared to those with CR alleles of the same uninterrupted CAG repeat lengths, suggesting that LI and DI alleles confer additional effects. Thus, we attempted to determine the levels of additional effects of LI and DI that were not explained by the uninterrupted CAG size by taking a mathematical approach that is similar to least square approximation. Briefly, we predicted age-at-onset of HD subject carrying LI or DI alleles using our CAG-onset regression model for CR ^19^, and subsequently calculated the residual by subtracting predicted onset from observed onset age. We then calculated the sum of square (SS) for LI and DI carriers based on the participant’s true uninterrupted CAG repeat length using the following formula.

Sum of squares (SS) = ∑ (observed age-at-onset − predicted age-at-onset age) ^2^

Subsequently, we gradually increased and decreased the CAG repeat length for LI and DI allele carriers, respectively, and calculated SS again to identify CAG repeat size that generated the smallest SS. The differences between true CAG and CAG repeat length that produced the smallest SS were considered as additional effects of LI and DI alleles on age-at-onset.

### HEK293 cell culture, gRNA cloning, and transfection

HEK293 cells (https://www.atcc.org/products/crl-1573) were maintained in DMEM containing L-glutamine supplemented with 10% (v/v) FBS and 1% Penicillin-Streptomycin (10,000U/ml). Cells were maintained at 37 °C and 5% CO2. TrypLE Express (Life Technologies) was used to detach cells for sub-culture. PX552 vector (addgene #60958) was digested using SapI (Thermo) and purified by gel purification (QIAquick Gel Extraction Kit). A pair of oligos for each sgRNA were phosphorylated (T4 Polynucleotide Kinase, Thermo) and annealed by incubating at 37 °C for 30 min, 95 °C for 5 min, and ramping to 4 °C. Annealed oligos were diluted (1:50) and ligated into the digested PX552 vector (T7 ligase, Enzymatic) and incubated at room temperature for 15 min. Then, transformation was performed (One Shot Stbl3, Invitrogen). The inserted gRNA sequences were confirmed by Sanger sequencing. For transfection, cells were seeded in 6-well plates at approximately 65% confluence and treated with 1.66ug of CBEs and 0.7ug of gRNA plasmids on the following day using Lipofectamine 3000 (Invitrogen) according to the manufacturer’s protocol. Three days after transfection, cells were harvested for molecular analyses. Genomic DNA was extracted using DNeasy Blood & Tissue kit (QIAGEN). AccuPrime GC-Rich DNA Polymerase (Invitrogen) was used to amplify a region containing the CAG repeat (35 cycles). PCR product was purified by PCR QIAquick PCR Purification Kit (QIAGEN) and subjected to MiSeq (Center for Computational and Integrative Biology DNA Core, Massachusetts General Hospital) and/or Sanger sequencing analysis (Center for Genomic Medicine Genomics Core, Massachusetts General Hospital). Primers for MiSeq sequencing were ATGAAGGCCTTCGAGTCCC and GGCTGAGGAAGCTGAGGA; primers for Sanger sequencing analysis were CAAGATGGACGGCCGCTCAG and GCAGCGGGCCCAAACTCA.

### MiSeq data analysis to determine indels and conversion types

Sequence data from the MiSeq sequencing were subject to quality control (QC) by removing sequence reads 1) with mean base Phred quality score smaller than 20, 2) showing the difference between forward and reverse read pair, 3) containing fewer than 6 CAGs, or 4) not involving the full primer sequences. QC-passed data revealed that HEK293 cells carry 16/17 CAG canonical repeats and therefore are expected to produce 18/19 polyglutamine segments. For QC-passed sequence reads, we determined the proportion of sequence reads containing indels, revealing most indels were sequencing errors. Subsequently, we focused on sequence reads without indels to determine the types of conversion. For each sequence read (not including CAA-CAG interruption), we counted sequence reads containing CAA, CAC, CAG, CCG, CGG, CTG, AAG, GAG, and TAG trinucleotide to determine the types and levels of conversion.

### MiSeq data analysis of HEK293 cells treated with BE strategies to determine the sites of conversion

Sequence analysis revealed that BE strategies using CBEs produced mostly CAG-to-CAA conversion. CAG-to-TAG conversion was detected in all samples regardless of BE strategies, suggesting that this type of conversion is also due to amplification/sequencing errors. Therefore, we focused on sequence reads of 16/17 CAG repeats containing only CAG or CAA to determine the sites of CAG-to-CAA conversion. Briefly, we recorded the sites of CAG-to-CAA conversion for each sequence read and summed the number of conversions at a given site for a given sample. Therefore, 30% CAG-to-CAA conversion at the second CAG means 70% and 30% of all sequence reads contain CAG and CAA at the second CAG position, respectively.

### Quantification of duplicated interruption and multiple conversion

The proportion of duplicated interruptions was determined from HEK293 cells treated with different combinations of CBEs and gRNAs. Briefly, we counted sequence reads containing duplicated interruption and divided them by the number of all QC-passed sequence reads to calculate the proportion of DI alleles. Similarly, we calculated the proportion of sequence reads containing duplicated interruption and CAG-to-CAA conversions at other sites. We also determined the levels of multiple CAG-to-CAA conversion for each BE strategy. For each sequence read in a sample, we counted the number of CAG-to-CAA conversions regardless of their locations to generate a distribution of numbers of multiple conversion for each sample. Since we counted the conversions regardless of their positions, multiple conversions do not necessarily mean consecutive conversions.

### Determination of the transfection efficiency

To determine the effects of transfection efficiency on patterns of base editing, we transfected HEK293 cells with gRNA 2 with combinations of different base editors and performed cell staining. Transfected HEK293 cells were fixed with paraformaldehyde (4%) and permeabilized with Triton X-100 (0.5%). Then, cells were stained with DAPI (0.5uM) and incubated for 30 min before being washed with PBS. The eGFP (enhanced green fluorescent protein) and DAPI (4’,6-diamidino-2-phenylindole) images from eight areas in each well were captured using a fluorescence inverted microscope (Nikon Eclipse TE2000-U). The ImageJ analysis program was used to measure the size of a single cell expressing eGFP; we randomly selected 20 cells for each image and averaged their sizes to be used as a reference. We counted the number of pixels covered by eGFP-positive signals, and subsequently divided by the average cell size to obtain the number of eGFP positive cells in each image. This was repeated with the DAPI staining images. The percent transfected was calculated by dividing the number of eGFP-positive cells by that of DAPI-positive cells multiplied by 100.

### Base editing in HD patient-derived iPSC and differentiated neurons

An iPSC line carrying adult-onset CAG repeats (42 CAG) was derived from a lymphoblastoid cell line in our internal collection by the Harvard Stem Cell Institute iPS Core Facility (http://ipscore.hsci.harvard.edu/) ^35; 36^. HD iPSC cells were dissociated into single cells with Accutase (STEMCELL Tech) and plated on matrigel-coated 24-well plate in mTeSR plus media containing CloneR (STEMCELL Tech) to increase cell viability. The following day, cells at 60∼70% confluence were transfected with 1.8ug of BE4max and 0.6ug of gRNA plasmids using Lipofectamine STEM (Invitrogen) according to the manufacturer’s protocol. Cells were incubated at 37 °C and 5% CO2 for 5 days for sequencing analysis.

The same iPSC line was differentiated into neurons using a previously described method ^37^. Briefly, the iPSC line was plated on growth factor reduced Matrigel (Corning) in mTeSR Plus media (STEMCELL Technologies). When cells reached ∼ 80% confluence, differentiation was initiated by switching to DMEM-F12/Neurobasal media (2:1) supplemented with N2 and retinol-free B27 (N2B27 RA−; Gibco). For the first ten days, cells were supplemented with SB431542 (10 µM; Tocris), LDN-193189 (100 nM; StemGene), and dorsomorphin (200 nM; Tocris). SB431542 was removed from the media on day 5. Cells were maintained in N2B27 RA− supplemented with activin A (25 ng/ml; R&D) on day 9. On day 22, cells were split using Accutase (STEMCELL Technologies) and seeded on a poly-D-lysine/laminin plate with N2B27 media supplemented with BDNF and GDNF (10 ng/ml each; Peprotech). Media were changed the next day to facilitate neuronal maturation and survival. Cells were fed with new media every two days. For neuronal marker staining, cells were fixed, permeabilized, and blocked using the Image-iT Fix-Perm kit (Invitrogen). Subsequently, cells were stained by Anti-TUBB3 (tubline beta 3; Biolegend Inc, Cat# 801202) in a blocking solution overnight at 4°. Then, cells were washed with PBS three times for 5 minutes, followed by incubation with Alexa Flour 594 secondary antibodies (Invitrogen) for 1 hour. Finally, cells were washed with PBS three times for 5 minutes and mounted with Vectorshield mounting medium with DAPI (Vector Laboratories). Images were captured by the Leica fluorescence microscope. Differentiated neurons were transfected with 1.8ug of BE4max and 0.6ug of gRNA plasmids using Lipofectamine 3000 according to the manufacturer’s protocol. Cells were incubated at 37 °C and 5% CO2 for 7 days for sequencing analysis.

### Off-target prediction and experimental validation

Potential off-targets were predicted by the Off-Spotter (https://cm.jefferson.edu/Off-Spotter/) for 8 BE strategies using the gRNA sequences. We allowed a maximum of 4 mismatches to identify potential off-targets that are flanked by the NGG PAM. Given decreased single base specificity at the PAM-proximal sites in the CRISPR-Cas9 genome engineering ^38^ and the abundance of CAG repeat carrying genes in the human genome, many of our gRNAs (except gRNAs 1 and 2) are predicted to hybridize with many CAG repeat sequences in the genome, generating increased numbers of predicted off-targets. Thus, we performed experimental validation of 1) predicted off-targets for BE strategies 1 and 2 (described here) and 2) genes that cause polyglutamine disorders. For the experimental validation of predicted off-targets, we analyzed HEK293 DNA samples that were used for MiSeq analysis. Briefly, we focused on predicted off-targets in the protein-coding genes for gRNAs 1 and 2 with two mismatches. One and four potential off-targets were predicted for gRNAs 1 (*MINK1*) and 2 (*PINK1, ZNF704, WBP1L, C20orf112*), respectively. We amplified predicted off-target sites of gRNAs 1 and 2 (35 cycles) using the following primers:

*MINK1*, AGCATGCCTACCTCAAGTCC and CTGGTTTGTCAGCGGGATTC;

*PINK1*, CTGTACCCTGCGCCAGTA and GGATGTTGTCGGATTTCAGGT;

*ZNF704*, GGACGGGTTGGACTGGTC and GGGTCCTGGCACTGACTGTG;

*WBP1L*, CCGACCTCCAACTCCTCCC and GCTGCTCTGTGCCCCCTG; and

*C20orf112*, GATCTCCGTGGGGCTGAG and CCTACTTCCCTCTCCACAGG.

Amplified DNA samples were analyzed by MiSeq sequencing.

### Experimental validation of off-targets in genes causing polyglutamine diseases

Similarly, we amplified genomic regions (35 cycles) containing CAG repeat regions in the genes causing polyglutamine diseases using the following primers:

ATXN1, CCTGCTGAGGTGCTGCTG and CAACATGGGCAGTCTGAGC;

*ATXN2*, CGGGCTTGCGGACATTGG and GTGCGAGCCGGTGTATGG;

*ATXN3*, GAATGGTGAGCAGGCCTTAC and TTCAGACAGCAGCAAAAGCA;

*CACNA1A*, CCTGGGTACCTCCGAGGGC and ACGTGTCCTATTCCCCTGTG;

*ATXN7*, GAAAGAATGTCGGAGCGGG and CTTCAGGACTGGGCAGAGG;

*TBP*, AAGAGCAACAAAGGCAGCAG and AGCTGCCACTGCCTGTTG;

*ATN1*, CCAGTCTCAACACATCACCAT and AGTGGGTGGGGAAATGCTC; and

*AR*, CTCCCGGCGCCAGTTTGCTG and GAACCATCCTCACCCTGCTG.

Sequencing data analysis was focused on calculating the proportion of sequence reads that contain the CAG-to-CAA conversions.

### RNAseq analysis

To determine the molecular consequences of candidate BE strategies, we performed RNAseq analysis. We transfected HEK293 cells with BE4max+empty vector, BE4max+gRNA 1, or BE4max+gRNA2 for 72hours. Subsequently, genomic DNA for MiSeq analysis and cell pellets for RNAseq analysis were generated from replica plates Genome-wide RNAseq analysis (Tru-Seq strand specific large insert RNA sequencing) was performed by the Broad Institute. Sequence data were processed by STAR aligner ^39^ as part of the Broad Institute’s standard RNAseq analysis pipeline. For differential gene expression (DGE) analysis, we used transcripts per million (TPM) data computed by the TPMCalculator (https://github.com/ncbi/TPMCalculator) ^40^. Expression levels in approximately 19,000 protein-coding genes based on Ensembl (ftp://ftp.ensembl.org/pub/release-75/gtf/homo_sapiens/) were normalized. The DGE analysis was performed by the generalized linear model using a library of “glm” in R package v3.3.1 (https://www.r-project.org/) after adjustment for two principal components based on RNAseq data, followed by multiple test correction using a FDR method. A multiple test corrected p-value less than 0.05 was considered statistically significant.

### Generation and validation of HEK293-51 CAG cells carrying an expanded CAG repeat

HD patient-derived iPSC and neurons showed modest conversion efficiencies, making it technically difficult to characterize molecular consequences of CAG-to-CAA conversion strategies. Thus, we generated HEK293 cells carrying an expanded repeat by replacing one of the non-expanded *HTT* CAG repeats with a 51 CAG repeat. Briefly, we cloned a gRNA (CAGAGCGCAGAGAATGCGCG) into the PX459 vector (Addgene# 62988) to express SpCas9 and gRNA for CRISPR-Cas9 targeting at the *HTT* CAG repeat region. The donor template for homologous recombination was generated by PCR amplification of a human DNA sample carrying 51 CAG repeat into the pCR-Blunt II TOPO plasmid (Invitrogen). Subsequently, HEK293 cells were transfected with PX459 and pCR-Blunt II TOPO plasmids by Lipofectamine 3000 (Invitrogen) for 72 hr. Subsequently, cells were treated with G-418 (Gibco) for 21 days, and surviving cells were re-plated onto 100 cm dishes. After 10 days, visible colonies were picked and maintained separately. Single cell clonal lines were validated by PCR analysis using AccuPrime GC-Rich DNA Polymerase and primer set (ATGAAGGCCTTCGAGTCCC and GGCTGAGGAAGCTGAGGA). The PCR conditions were initial denaturation (95 °C, 3 min), 30 cycles of denaturation (95 °C, 30 sec), annealing (55 °C, 30 sec), extension (72 °C, 40 sec), and final extension (72 °C, 10 min). The PCR products were resolved on a 1.5% agarose gel containing GelRed (Biotium) and visualized under UV light to distinguish expanded from non-expanded CAG repeats. We also performed RT-PCR and immunoblot analysis to confirm the correct integration of the expanded CAG repeat. Briefly, 1 μg of total RNA from the targeted clonal line was subjected to reverse transcription with SuperScript IV Reverse Transcriptase (Invitrogen) according to the manufacturer’s instructions followed by PCR analysis using a primer set (ATGAAGGCCTTCGAGTCCC and GGCTGAGGAAGCTGAGGA). For HTT immunoblot analysis, cells were lysed with RIPA Lysis/Extraction Buffer (Thermo) supplemented Halt Protease and Phosphatase Inhibitor Cocktail (Thermo). Whole cell lysate was then separated on NuPAGE 3 to 8%, Tris-Acetate gel (Invitrogen) and transferred to a polyvinylidene fluoride membrane. The membrane was blocked with 5% nonfat dry milk in Tris-buffered saline for 1 h and incubated with primary antibodies for HTT (MAB2166, Sigma-Aldrich) for 12 h at 4 °C. The membrane was washed for 1 h, and blots were incubated with a peroxidase-conjugated secondary antibody for 1 h then washed for 1 h. The bands were visualized by enhanced chemiluminescence (Thermo). Similar to HEK293 cells, HEK293-51CAG cells were treated with BE4max and candidate gRNAs (i.e., gRNA 1 and gRNA 2) to determine the levels of CAG-to-CAA conversion and the total HTT protein levels. For gRNA 1, we determined the levels of in-frame insertion and deletion right after treatment using methods previously described (Lee 2015).

### AAV treatment for a candidate BE strategy and CAG repeat instability in mice

For AAV injection experiments, we used split-intein base editor (v5 AAV) ^34^. Forward and reverse oligos (CACCGCTGCTGCTGCTGCTGCTGGA and AAACTCCAGCAGCAGCAGCAGC) (IDT) for gRNA 2 were cloned into the BSmBI site of pCbh_v5 AAV-CBE C-terminal (Addgene, # 137176) and pCbh_v5 AAV-CBE N-terminal (Addgene, # 137175). Cloned vectors were validated by Sanger sequencing, and subsequently packed into AAV9 serotype by UMass Viral Vector Core. HttQ111 HD knock-in mice ^41^ were maintained on an FVB/N background ^42^; AAV9 injections were performed in heterozygous HttQ111/+ mice at 6∼11 week. Animal husbandry was performed under controlled temperature and light/dark cycles. After anesthesia was induced using isoflurane, an insulin syringe was inserted into the medial canthus with the bevel of the needle facing down from the eyeball, advanced until the needle tip was at the base of the eye. We injected HD knock-in mice with AAV9 mix (200 μl containing C-terminal and N-terminal split-intein base editor, 1 x 10^12^ vg for each) (experimental group) or PBS (200 μl, control group) by retro-orbital (RO) injection. Ten weeks later, liver and tail samples were collected for instability analysis ^43; 44^. Briefly, DNA samples were amplified using primer set (6’FAM-ATGAAGGCC TTCGAGTCCCTCAAGTCCTTC and GGCGGCTGAGGAAGCTGAGGA) and analyzed by ABI3730 to determine the sizes of fragments. Quantification of repeat expansion was based on the expansion index method that we developed previously. The expansion index method robustly quantifies the levels of repeat instability by eliminating potential noise in the fragment analysis results based on the relative peak height threshold ^43; 44^. To quantify expansion index in control and mice treated with BE, we applied 10% threshold, and expansion index was calculated based on the highest peak in the tail DNA.

### Repeat instability in HD knock-in mice carrying interrupted CAG repeat

To determine the maximal effects of CAG-to-CAA interruption, we analyzed HD knock-in mice carrying interrupted CAG repeat (namely interrupted repeat mice; https://www.jax.org/strain/027418) to HD knock-in mice carrying uninterrupted repeat (namely, pure repeat mice; https://www.jax.org/strain/027417). Repeat in the interrupted repeat mice and pure repeat mice comprises 21 copies of [CAGCAACAGCAACAA] and 105 copies of [CAG], respectively. Both mouse lines were expected to produce huntingtin protein with 105 polyglutamine. Repeat instability in these mice were determined (5 months) by the fragment analysis as described previously ^44^.

### Statistical analysis and software

Statistical analysis of RNAseq data was performed using generalized linear regression analysis. Multiple test correction was performed using false discovery rate using R 3.5.3 ^45^. R 3.5.3 was also used to produce plots.

## Results

### Effects of CAG-CAG codon doublet on age-at-onset in HD patients

Previously, we and others reported that most HD subjects carry canonical repeats (CR) comprising an uninterrupted expanded CAG repeat followed by CAA-CAG ^24; 27; 30^. Although infrequent, uninterrupted CAG repeats followed by 1) no CAA-CAG (LI; 0.23% in our previous GWA data) and 2) two CAA-CAG codon doublets (DI; 0.76% in our previous GWA data) also exist (S. Figure 1). In HD subjects carrying LI alleles, the length of the CAG repeat and polyglutamine segment are identical. However, the polyglutamine length is greater by 2 and 4, respectively, compared to the CAG repeats in CR and DI alleles (S. Figure 1). Since CR, LI, and DI alleles with the same uninterrupted CAG repeat lengths have different polyglutamine sizes, they have provided a powerful tool to investigate the relative importance of the CAG repeat in DNA vs. polyglutamine in protein in determining onset age. For example, if polyglutamine length played an important role in determining age-at-onset, onset of LI and DI allele carriers, who respectively have 2 fewer and 2 more glutamines compared to CR allele carriers, would be significantly later and earlier compared to CR allele carriers with the same uninterrupted CAG repeats (S. Figures 2A and 2B). In stark contrast to these predictions, the onset ages of LI or DI allele carriers are best explained by their respective CAG repeat sizes, not polyglutamine length (S. Figures 2C and 2D). Furthermore, age-at-onset of DI allele carriers is significantly delayed compared to that of LI allele carriers with the same uninterrupted CAG repeat size even though DI alleles encode 4 more glutamines than LI alleles (Student t-test p-value, 1.007E-12) (S. Figure 2D). Together, the data indicate that age-at-onset in HD is determined primarily by the uninterrupted length of the CAG repeat, but there may also be additional effects of different CAA-interruption structures since the CAG repeat length does not fully explain age-at-onset in LI and DI allele carriers (S. Figure 2D) ^46^. Therefore, we performed least square approximation to calculate the magnitudes of the additional effects of LI and DI alleles on age-at-onset. Briefly, we varied the individual CAG repeat length to identify the repeat size that best explains the observed age-at-onset of carriers of these LI and DI alleles relative to CR alleles. The age-at-onset of the LI allele carriers is best explained when 3 CAGs are added to the true CAG repeat length (Figures 1A and 1B; S. Figure 3D) while the DI allele carriers behave with respect to age-at-onset as if they have one less CAG than their true CAG repeat length (Figures 1B and 1C; S. Figure 3F). These data suggest that switching a CR allele to a DI allele would delay onset by 1) shortening the uninterrupted CAG repeat by two CAG repeats and 2) conferring an additional effect comparable to a reduction in length of one CAG. For example, if a DI allele were generated from a CR allele with 43 uninterrupted CAGs by converting the 42nd CAG to CAA using base editing strategies, the age-at-onset is predicted to be delayed by approximately 12 years (Figure 1D), illustrating the robustness of therapeutic base editing strategies.

**Figure 2.**
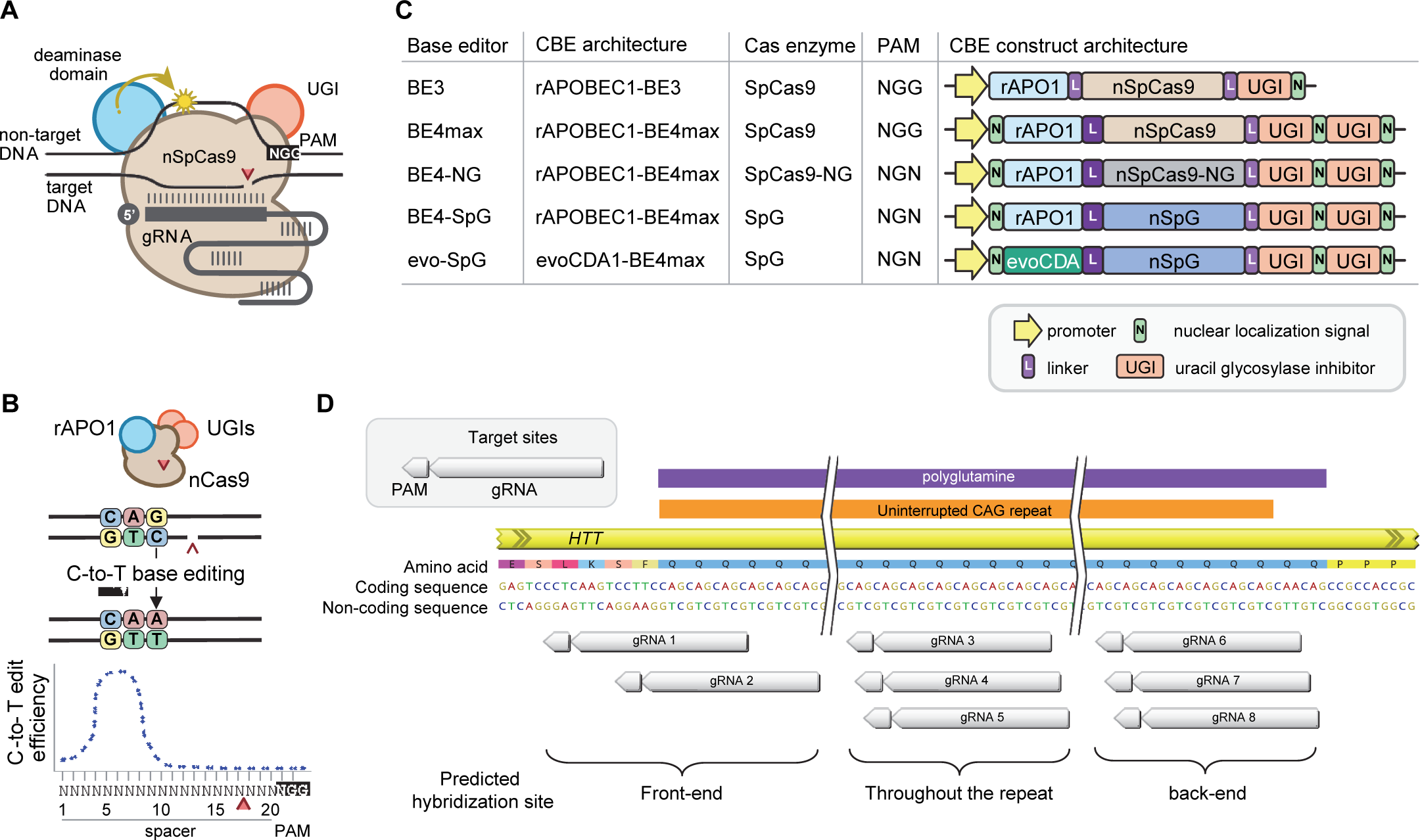
Cytosine base editors and gRNAs for CAG-to-CAA conversion in HD. (A) Constituents of base editing are displayed. (B) Schematic of cytosine base editors (CBEs) that can generate C-to-T edits within a finite edit window at a fixed distance from the PAM. (C) CBE variants described in the literature and used in this study (lower 4) are shown, including the evoCDA1- based SpG CBE that should function more efficiently in GC nucleotide contexts. Protospacer-adjacent motif (PAM), guide RNA (gRNA), uracil glycosylase inhibitor (UGI), rat APOBEC1 deaminase domain (rAPO1), evolved CDA1 cytosine deaminase domain (evoCDA). (D) The target region, gRNAs, and expected hybridization sites of the 8 gRNAs are shown.

**Figure 3.**
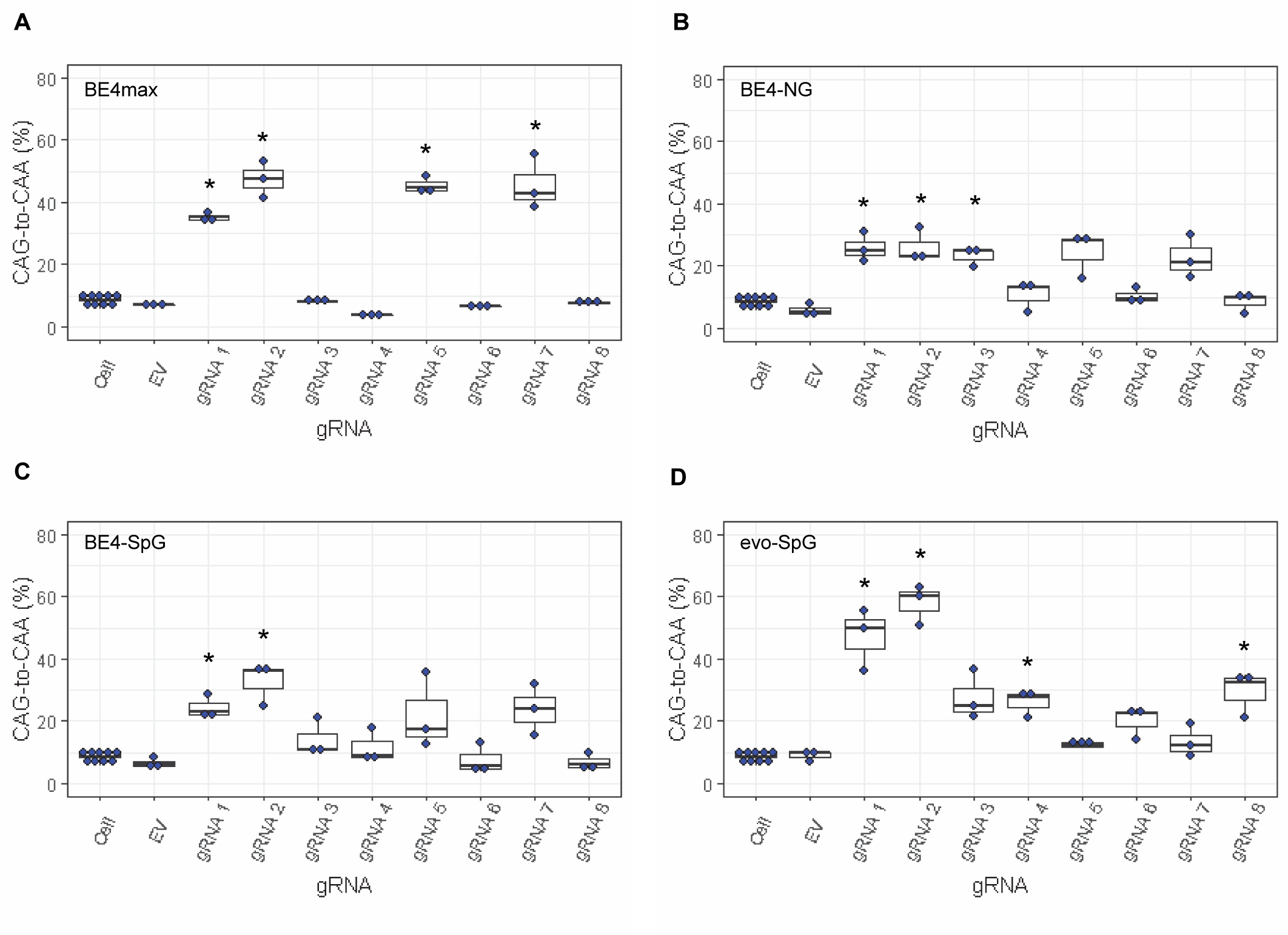
Levels of CAG-to-CAA conversion by BE strategies. Only CAG-to-CAA conversion showed significantly increased levels over the baseline sequencing errors. Thus, we calculated the percentage of CAA in the cells that were treated with a combination of cytosine base editors (A, BE4max; B, BE4-NG; C, BE4-SpG; and D, evo-SpG) and gRNAs. HEK293 cells without any treatment (i.e., Cell) were combined (n=8) and plotted for each base editor. EV represents HEK293 cells treated with a base editor and empty vector for gRNA. *, significant by Bonferroni corrected p-value < 0.05 (8 tests for each base editor).

### Cytosine base editors and gRNAs to convert CAG to CAA in the HTT CAG repeat

Recent advancements in genome editing technologies have led to the development of CBEs that are capable of efficient C-to-T conversion (Figure 2A) ^31; 32; 47–49^. In principle, CR can be converted to DI if CBEs target the non-coding strand of the *HTT* CAG repeat (Figure 2B). In this study, we tested 4 CBEs comprised of various cytosine deaminases and SpCas9 enzymes with different protospacer-adjacent motif (PAM) specificities to explore the feasibility of CAG-to-CAA conversion as a putative treatment for HD. BE4 is the fourth-generation base editor which was engineered from BE3 to increase the editing efficiencies and decrease the frequency of undesired by-products (Figure 2C) ^47^. BE4 exhibited high levels of C-to-T editing activity on the target sites harboring NGG PAMs ^47^. The activity window of BE4 is position 4-8, counting from the PAM distal end of the spacer (where the PAM is positions 21-23) (Figure 2B) ^48^. We tested the BE4max (Addgene #112093) in this study, which is a codon optimized version of BE4 with improved nuclear localization ^48^. Due to the sparsity and lack of NGG PAM sites near and within the CAG repeat, respectively, CAG-to-CAA conversion using BE4 was expected to be somewhat limited. Therefore, we also explored engineered CBEs containing SpCas9 variants that target an expanded range of PAM sequences, including SpCas9-NG ^50^ and SpG ^51^ (Figure 2C). Since these variants are capable of targeting sites with NGN PAMs, they might permit higher density targeting near or within the CAG repeat. The nucleotide preceding the target cytosine also affects the C-to-T conversion efficiency in CBEs, especially when a G precedes the C ^31; 52; 53^. Thus, engineered deaminase domains have been explored to improve C-to-T conversion in the GC-contexts ^32; 49^. For instance, an evolved CDA1-based BE4max variant (evoCDA1) showed substantially higher editing on GC targets ^49^, which is relevant to the nucleotide context on the non-coding strand of the *HTT* CAG repeat (CT<u>GC</u>TG). Therefore, we explored the use of canonical BE4max-SpCas9, BE4max-SpCas-NG, BE4max-SpG, and evoCDA1-BE4max-SpG (henceforth referred to as BE4max, BE4-NG, BE4-SpG, and evo-SpG, respectively) (Figure 2C).

To achieve CAG-to-CAA conversion in the *HTT* CAG repeat, we designed 3 groups of gRNAs (S. Table 1) based on the sites of predicted hybridization (Figure 2D). Aiming at converting CAGs at the front-end of the repeat, gRNAs 1 and 2 were designed to hybridize with a region involving the upstream of the repeat and conventional NGG PAMs. The gRNAs 1 and 2 contain 10 and 2 non-CAG bases at the PAM-proximal ends, respectively (S. Table 1; S. Figure 4). Considering the activity window of the BE4 (i.e., 13th-17th nucleotide from the PAM) (S. Figure 4, green boxes in the gRNAs), BE4max-gRNAs 1 and BE4max-gRNA 2 were predicted to convert the 1st/2nd and 4th/5th CAG to CAA, respectively (S. Figure 4; sequences with green highlight). The gRNAs 3, 4, and 5 comprised the CAG repeat sequence (S. Table 1) and therefore, were predicted to hybridize throughout the *HTT* CAG repeat and potentially other CAG repeat-containing genes (S. Figure 4). The gRNAs 3, 4, and 5 were predicted to utilize NAA/NTG, NGA/NCT, and NGG/NGC PAMs, respectively (S. Figure 4). Lastly, gRNAs 6, 7, and 8 were designed to convert CAGs at the back-end of the repeat (S. Table 1). Available PAM sites for these gRNAs are NCT, NGC, and NTG (S. Figure 4). Considering the predicted gRNA-target hybridization sites and conversion windows, these three gRNAs might generate the duplicated interruption that is found in HD patients. (S. Figure 4).

**Figure 4.**
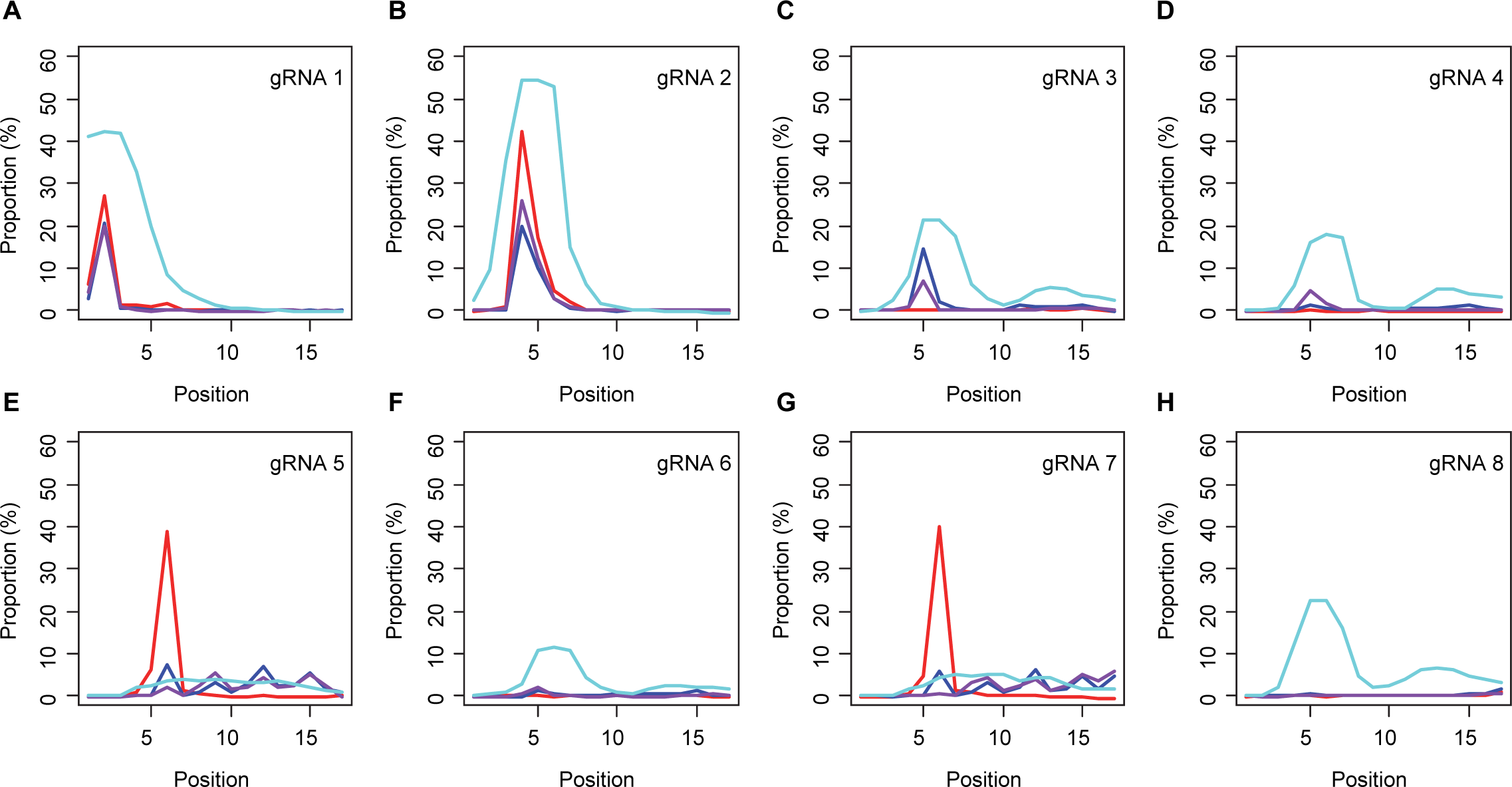
Sites of CAG-to-CAA conversion by BE strategies. We calculated the percentage of sequence reads containing CAA at specific sites relative to all sequence reads. For example, 27.7% conversion at the 2nd CAG by BE4max-gRNA 1 (top left panel, red) means 27.7 % of all sequence reads from the 16 or 17 CAG alleles have CAA at the 2nd CAG. X-axis and y-axis represent the position of the CAG and percent conversion. Each panel represents a tested gRNA. Plots were based on the mean of 3 independent transfection experiments in HEK293 cells after subtracting corresponding empty vector (EV)-treated cell data. Red, blue, purple, and cyan traces represent BE4max, BE4-NG, BE4-SpG, and evo-SpG.

### Predominant CAG-to-CAA conversion without significant indels by BE strategies for HD

We then characterized 32 BE strategies (i.e., combinations of 4 CBEs and 8 gRNAs). We first determined whether BE strategies for HD produced indels. Since low base editing efficiencies might result in proportionally low levels of indels leading to an underestimation of their frequencies, we used HEK293 cells, which showed high levels of base editing efficiencies ^54; 55^. Our MiSeq sequence analyses revealed that HEK293 cells carry two CRs (16 and 17 CAGs) and showed approximately 10% of basal levels of indel (’Cell’ in S. Figure 5), which reflects errors due to the difficulty in sequencing the CAG repeat. Nevertheless, transfection of plasmids for BE strategies did not significantly increase the levels of indels compared to cells without any treatment (Cell) or cells treated with empty vector (EV) (S. Figure 5). The lack of significant indel formation was quite expected because the cytosine base editors that we tested use nickases (Figure 2C) ^47; 48^.

**Figure 5.**
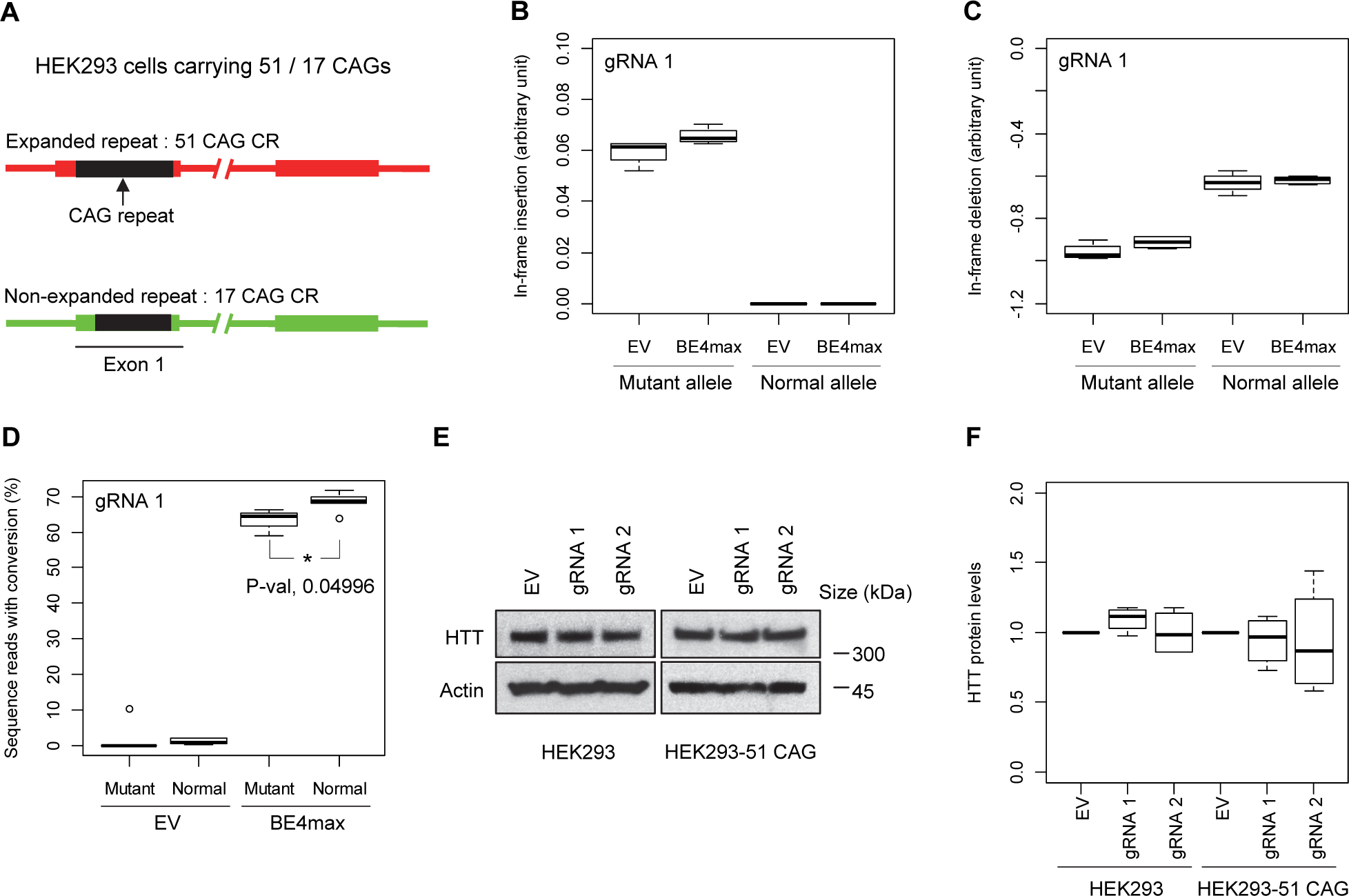
Allele specificity and molecular outcomes of candidate BE strategies. (A) To overcome the limitations of patient-derived iPSC and differentiated neurons, we developed HEK293 carrying an adult-onset CAG repeat by replacing one of the normal repeats with 51 canonical CAG (namely HEK293-51 CAG). Red and green bars represent respectively mutant and normal *HTT* in HEK293-51 CAG cells. (B and C) The HEK293-51 CAG cells were treated with BE4max-gRNA 1 and analyzed to determine the levels of in-frame insertion (B) and in-frame deletion (C) at the time of treatment. (D) The HEK293-51 CAG cells were treated with the gRNA 1 and analyzed by MiSeq to determine the levels of allele specificity. Conversion efficiency on the Y-axis indicates the percentage of sequence reads containing the CAG-to-CAA conversion at the target site. * represents uncorrected p-value < 0.05 by Student t-test. (E) Original HEK293 cells and HEK293-51 CAG cells were treated with empty vector (EV), or candidate BE strategies (BE4max-gRNA 1 and BE4max-gRNA 2) and subjected to immunoblot analysis; representative blot is shown in panel E. (F) Four independent experiments were performed, and we performed one-sample t-test to determine whether BE-treated cells show different total HTT protein levels compared to EV-treated cells. Nothing was significant by p-value < 0.05.

Since most sequence reads containing indels might be sequencing errors, we focused on sequence reads without indels to determine the types of base conversions. HEK293 cells without any treatment (Cell) or cells treated with empty vector (EV) showed low but detectable levels of CAG-to-CAA and CAG-to-TAG conversions (S. Table 2; S. Figure 6), also reflecting sequencing errors. However, the levels of CAG-to-CAA conversion were significantly increased over baseline sequencing errors in cells treated with some BE strategies (S. Table 2). For example, BE4max in combination with gRNAs 1, 2, 5, and 7 resulted in efficient CAG-to-CAA conversion (Figures 3A). Given the availability of the NGG PAMs (S. Figure 4), robust CAG-to-CAA conversion by gRNAs 1, 2, and 5 was somewhat anticipated for BE4max. However, high levels of CAG-to-CAA conversion by the BE4max-gRNA 7 combination (Figure 3A; S. Figure 6A) were unexpected because the anticipated hybridization site does not provide the NGG PAM (S. Figure 4) that is required for the optimal activity of BE4max. The BE4-NG robustly produced CAG-to-CAA conversions with gRNAs 1, 2, and 3; although not significant, gRNAs 5 and 7 also generated high levels of CAG-to-CAA conversions (Figure 3B; S. Figure 6B). BE4-SpG with the combinations with gRNAs 1 and 2 resulted in significant levels of CAG-to-CAA conversions (Figure 3C; S. Figure 6C). Overall, the CAG-to-CAA conversion was higher in evo-SpG compared to other base editors; gRNAs 1, 2, 4, and 8 produced significant CAG-to-CAA conversions (Figure 3D; S. Figure 6D). These data indicated that our BE strategies primarily generated CAG-to-CAA conversion without significant indel formation. Patterns of conversions also indicated that sites with NGG PAMs (gRNAs 1, 2, and 5) permitted the highest levels of CAG-to-CAA conversion for BE4max and CBEs with relaxed PAM specificities.

**Figure 6.**
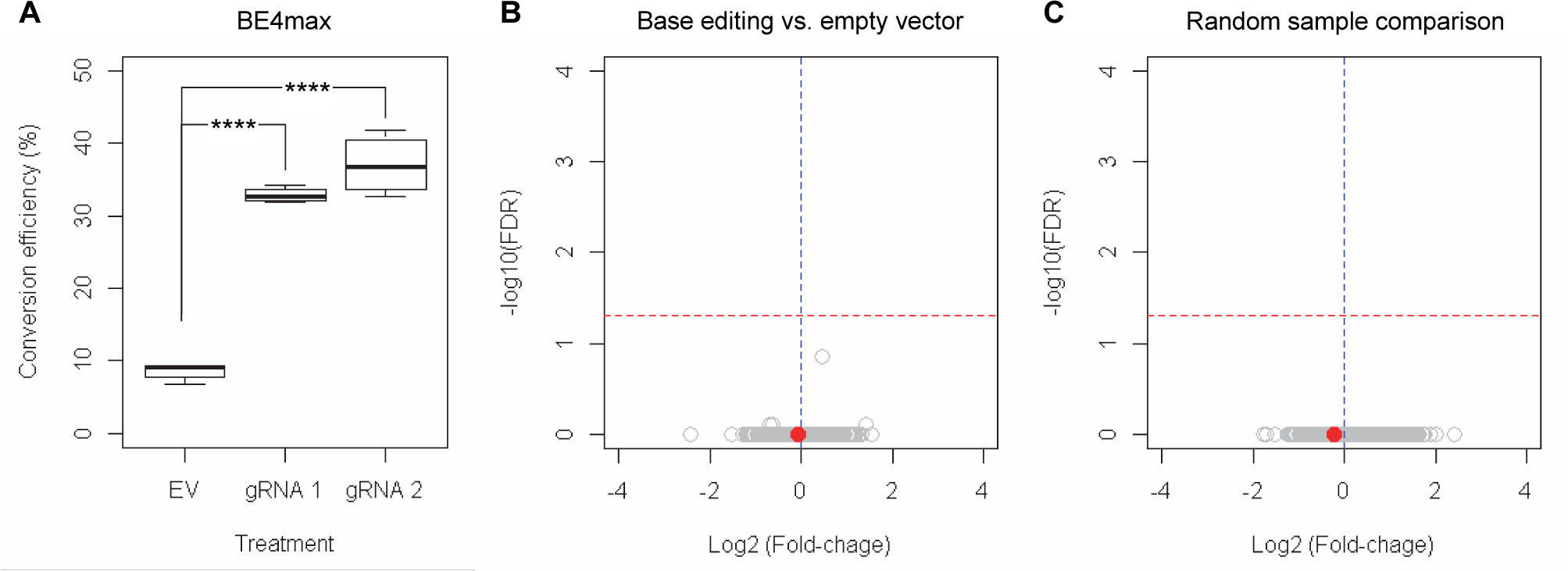
RNAseq analysis of BE strategies confirms the lack of transcriptome alternation. (A) HEK293 cells were treated with empty vector (EV) or candidate BE strategies such as BE4max-gRNA 1 (gRNA 1), and BE4max-gRNA 2 (gRNA 2) for RNAseq analysis. MiSeq analysis was also performed to judge the levels of CAG-to-CAA conversion. ****, p-value < 0.0001 by Student t-test (n=4). (B) Confirming the lack of significantly altered genes in BE4max-gRNA 1 or BE4max-gRNA 2, we compared all BE-treated samples (n=8) with all EV-treated samples (n=4) to increase the power in the RNAseq differential gene expression analysis. Each circle in the volcano plot represents a gene analyzed in the RNAseq; *HTT* is indicated by a filled red circle. A red horizontal line represents false discovery rate of 0.05, showing that none was significantly altered by candidate BE strategies. (C) We also compared two groups of randomly assigned samples (6 samples vs. 6 samples) to understand the shape of the volcano plot when there were no significant genes.

### Sites of CAG-to-CAA conversion

Subsequently, we determined conversion sites for different BE strategies. The patterns of conversion sites were similar for BE4max, BE4-NG, and BE4-SpG in gRNA 1, showing the most conversion at the second CAG with decreased levels of conversion at the first CAG (Figure 4A; S. Table 3). In contrast, evo-SpG-gRNA 1 combination showed higher editing efficiencies with the maximum conversion at the second CAG with comparable levels of conversions at the first and third CAGs (Figure 4A, cyan; S. Table 3). The gRNA 2 showed similar patterns as gRNA 1 except that conversion sites were shifted to the right; the highest conversion occurred at the 4th CAG by BE4max, BE4-NG, and BE4-SpG (Figure 4B; S. Table 3).

The gRNAs 3 and 4, which were designed to hybridize throughout the CAG repeat, did not generate CAG-to-CAA conversion in combination with BE4max (Figures 4C and 4D, red; S. Table 3) because of the lack of a NGG PAM. Although modest, BE4-NG and BE4-SpG converted the 5th CAG to CAA (Figures 4C and 4D; S. Table 3), potentially due to the possibility that NAA (gRNA 3) and NGA (gRNA 4) PAMs supported the base editing activity of BE4-NG and BE4-SpG. The gRNAs 3 and 4 produced higher levels of CAG-to-CAA conversion in evo-SpG again (Figures 4C and 4D, cyan), and interestingly, CAG-to-CAA conversions were not limited to the 5th CAG (S. Table 3). The gRNA 5 with BE4max efficiently converted the 6th CAG (Figure 4E, red), which was unexpected; conversions by other base editors were lower but widespread throughout the repeat (Figure 4E).

BE strategies designed to convert CAGs at the back-end of the repeat were tested using gRNA 6, 7, and 8. Although less robust, the patterns of conversion by gRNA 6 (Figure 4F) were similar to those of gRNA 4 (Figure 4D). Since only one nucleotide is different between gRNA 6 and gRNA 4, it appeared that gRNA 6 behaved like gRNA 4 despite one mismatch, favoring the NGA PAM instead of the less optimal NCT PAM (S. Figure 7A). The same explanation might account for the similar patterns of conversion sites for gRNA 7 (Figure 4G) and gRNA 5 (Figure 4E); efficient conversion at the 6th CAG by BE4max-gRNA 7 might be due to the interaction of gRNA 7 at the target site of gRNA 5 (with one mismatch) in favor of the NGG PAM (S. Figure 7B). The gRNA 8 generated CAG-to-CAA conversions only in evo-SpG. Although this group of gRNAs was designed to hybridize with the back-end of the CAG repeat, higher levels of conversion were observed at the front-end CAGs and throughout the repeat (Figures 4F-4H). These results suggest that one PAM-distal mismatch might be tolerated by base editors in favor of targets sites harboring more robust PAMs. Also, our data revealed that as expected, BE4max is highly dependent on the NGG PAM, resulting in CAG-to-CAA conversion at specific CAG sites, while evo-SpG is more efficient in conversion leading to broader targeting due to its relaxed PAM requirement.

**Figure 7.**
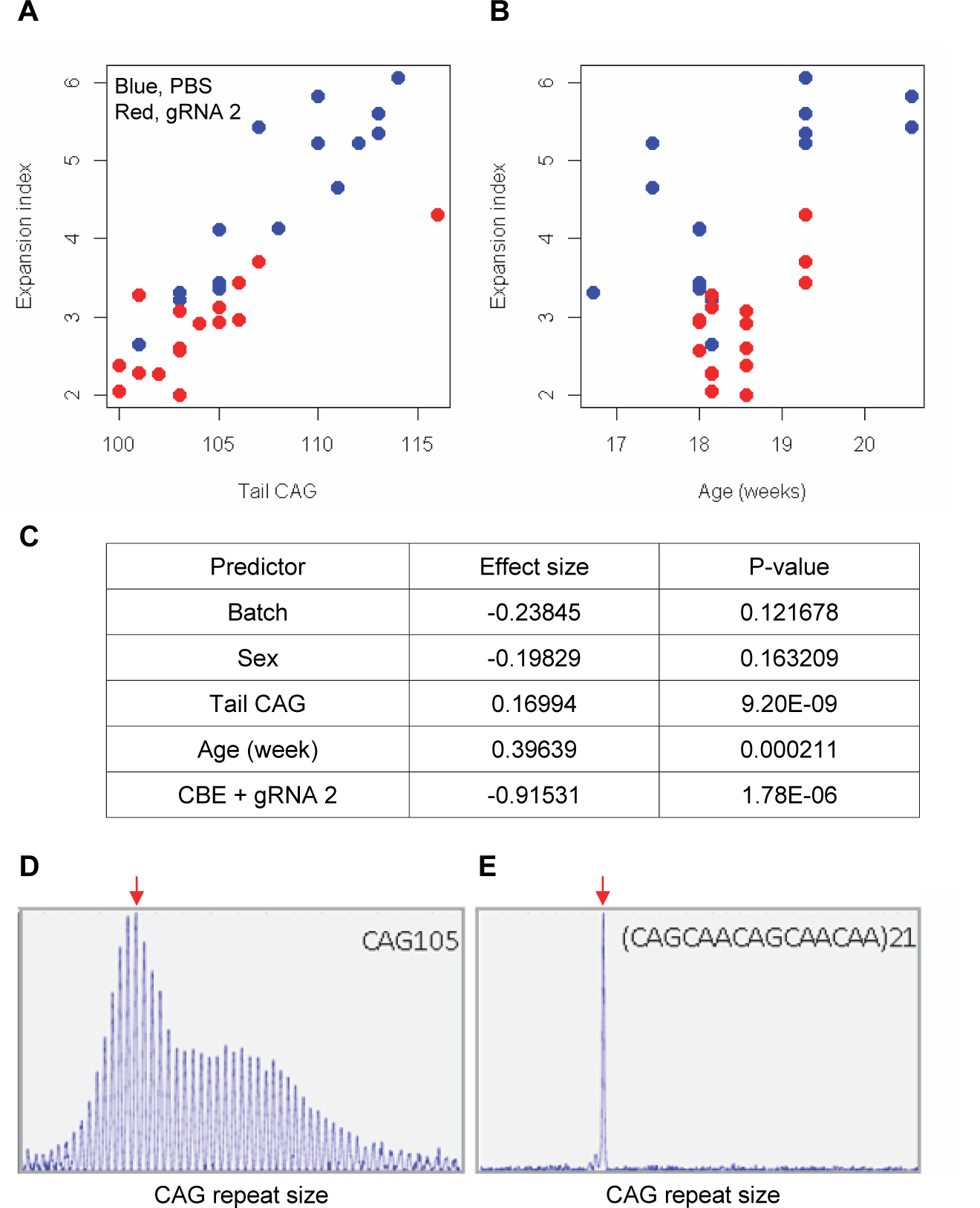
Impacts of CAA interruption on CAG repeat instability. (A - C) DNA samples (liver and tail) of BE-treated mice were analyzed to quantify somatic repeat expansion. We performed linear regression analysis to model the levels of repeat expansion as a function of treatment, CAG repeat in tail (A), age (B), and with other covariates (i.e., experimental batch, sex, tail CAG and age). Summary of the statistical analysis is summarized in the panel C. (D and E) To determine the maximal impacts of CAA interruption on the repeat expansion, HD knock-in mice carrying CAA interrupted repeats were analyzed. Liver samples of 105 uninterrupted CAG repeat (D) and interrupted repeat (E) were analyzed at 5 months. Representative fragment analysis is displayed. Red arrows indicate the modal alleles representing inherited CAG repeats; peaks at the right side of the modal peaks (red arrows) represent expanded repeats.

### Generation of duplicated interruption by BE strategies

Next, we determined the levels of duplicated interruption in the same HEK293 cell MiSeq data. As shown in S. Figure 8, BE4max and evo-SpG did not produce significant amounts of the DI that is found in humans (S. Figure 1). However, BE4-NG (S. Figure 8B) and BE4-SpG (S. Figure 8C) produced modest but significant levels of DI in combinations with gRNAs 5 and 7 (0.5%∼1% increase over the basal levels). Modest levels of DI alleles compared to conversions at other sites might be due to the lack of canonical PAMs (e.g., NGG) at the specific site (approximately 18 nucleotides upstream of CAA-CAG interruption). We also observed that gRNA 5 and 7 relatively increased the number of sequence reads containing both DI and CAG-to-CAA conversions at other sites (S. Figures 9B and 9C), indicating that CTG trinucleotides on the non-coding strand of the repeat contributed to modest but widespread CAG-to-CAA conversion throughout the repeat. Similarly, CAG-to-CAA conversion was not confined to specific sites in evo-SpG in combination with gRNAs 3-8 as DI alleles generated by these strategies also contained CAG-to-CAA conversions at other sites (S. Figure 9D). Since increased conversion efficiency in evo-SpG could not be explained by the transfection efficiency (S. Figure 10), these data indicate that evo-SpG has a significantly wider conversion window. In agreement with this, the most frequent number of conversions in a given sequence read by evo-SpG was greater than that of other base editors (S. Figure 11; S. Table 4).

### Evaluation of off-target effects

We then evaluated the levels of off-target conversions using Off-Spotter. As summarized in S. Table 5, gRNAs 1 and 2 showed relatively smaller numbers of predicted off-targets due to unique sequences near the PAMs. As expected, gRNAs that were designed to hybridize throughout the CAG repeat showed increased numbers of predicted off-targets. Similarly, gRNAs to convert CAG at the back-end of the repeat showed larger numbers of predicted off-targets, potentially due to the fact that unique sequences are distal to the PAMs. Subsequently, we performed two sets of follow-up off-target validations. For gRNAs 1 and 2, we experimentally evaluated predicted off-targets focusing on protein-encoding genes; one and four genes were predicted off-target sites for gRNA 1 and gRNA 2, respectively, and all showed low levels of conversion compared to on-target (S. Table 6). We also characterized the levels of off-target conversion in other CAG repeat-containing genes focusing on 8 polyglutamine disease genes (S. Tables 7). As predicted, gRNAs 1 and 2 showed low-level conversions in the CAG repeats of other polyglutamine disease genes in general (S. Table 8; S. Figure 12). In contrast, gRNAs 3- 8 produced variable but higher levels of conversion in some polyglutamine disease genes depending on the availability of preferred PAMs (S. Figure 12).

### Allele specificities and molecular outcomes of candidate BE strategies

Subsequently, we evaluate the levels of allele specificity of candidate BE strategies (BE4max-gRNA 1 and BE4max-gRNA 2) in patient-derived induced pluripotent stem cells (iPSC carrying 41 CAG CR) ^35; 36^ and differentiated neurons (S. Figure 13). As shown in S. Table 9, transfection of gRNA 1 and gRNA 2 produced modest CAG-to-CAA conversion on both mutant and normal *HTT* (approximately < 3%). Overall, low conversion efficiencies by transfection and transduction of AAV (adeno-associated virus; data not shown) represent difficulties in delivery in these cell types ^35; 56^, posing a challenge to determining the levels of allele specificity of BE strategies. To overcome these technical difficulties, we developed a HEK293 clonal line carrying an expanded *HTT* CAG repeat (Figure 5A) by replacing one of normal CAG repeats with a 51 CAG canonical repeat (namely HEK293-51 CAG) (S. Figure 14). A candidate BE strategy (i.e., BE4max-gRNA 1) did not increase the levels of in-frame insertion/deletion in the mutant or normal *HTT* repeat (Figures 5B and 5C). Subsequent analysis revealed that a candidate BE strategy BE4max-gRNA 1 produced high levels of CAG-to-CAA conversions on both expanded and non-expanded canonical repeats (Figure 5D). Although very modest, conversion was significantly higher in the non-expanded repeat (uncorrected p-value, 0.04996), which can be explained by slightly reduced conversion on the mutant *HTT* due to higher GC content in the expanded CAG repeat. However, the candidate BE strategies did not alter huntingtin protein levels (Figures 5E and 5F) at the time of treatment, supporting the safety of candidate BE strategies. We also performed RNAseq analysis to identify genes whose expression levels were altered by candidate BE strategies in HEK293 cells. Candidate strategies such as BE4max-gRNA 1 and BE4max-gRNA 2 produced significant on-target CAG-to-CAA conversions (Figure 6A), but the levels of *HTT* mRNA were not altered by either treatment (filled red circles in S. Figure 15 and Figure 6). In addition, RNAseq data analysis showed that neither BE strategy induced significant gene expression changes in any genes (false discovery rate, 0.05) (S. Figure 5). When comparing all HEK293 samples treated with either BE strategies (n=8) to those treated with EV (n=4), the shape of volcano plot mimicked random sample comparison (Figures 6B and 6C), implying the lack of impacts of candidate BE strategies on transcriptome.

### Effects of base conversion on the CAG repeat instability in vivo

The limited cargo capacity of AAV has been circumvented by the intein-split base editor, and the feasibility of BE strategies targeting non-repetitive sequences has been demonstrated in mouse models of human diseases ^34; 57; 58^. Taking advantage of the split BE system, we determined whether a candidate HD BE strategy could target the CAG repeat and result in a decrease in somatic repeat expansion, which was hypothesized to be the major disease driver ^20^. Since striatal and liver repeat instability share certain underlying mechanisms ^59–66^, and *in vivo* delivery might be more efficient in the liver compared to the brain ^34^, we used AAV9 to evaluate a candidate BE strategy in the liver. As expected, somatic CAG repeat expansion index in the liver of HD knock-in mice carrying around 110 CAGs showed a positive correlation with the inherited CAG repeat length (as represented in tail DNA) and the age of mice (Figures 7A and 7B) ^16; 23; 41; 44; 67; 68^. Unfortunately, we could not determine the sequence modification in treated mice by sequencing because of 1) very long CAG repeats in these mice, 2) modest levels of base conversion, and 3) high levels of errors when sequencing the CAG repeat (S. Figure 6). However, when the effects of the tail CAG repeat size and age of mice were corrected, retro-orbital injection of AAV9 for split CBE (v5 AAV-CBE) and gRNA 2 significantly decreased the levels of repeat expansion (Figure 7C; p-value, 1.78E-6). Nevertheless, the expansion index in treated and control mice was largely overlapping (Figure 7A), suggesting that the effects of BE treatment were very modest. We speculate that 1) insufficient dosage due to difficulty in producing high titer viral package for big cargo (i.e., 5KB) ^34; 69^, 2) limited delivery ^70^, and/or 3) difficulty in targeting the very long CAG repeat resulted in modest effects. Given those limitations, we also analyzed a mouse model containing interrupted repeat to determine the maximum effects of the interruption on the repeat expansion. HD knock-in mice carrying 105 interrupted CAG repeat (https://www.jax.org/strain/027418) showed complete loss of repeat expansion compared to 105 CAG uninterrupted repeat mice (https://www.jax.org/strain/027417) (Figures 7D and 7E), suggesting that CAA interruption could completely suppress the most important disease modifier (i.e., CAG repeat expansion).

## Discussion

Recent advances in genome engineering provide powerful tools to interrogate the relationships among genes, functions, and diseases. For example, CRISPR-Cas9-based editing approaches have revolutionized the investigation of individual genes of interest and also have begun to be applied to humans to treat diseases ^71–75^. Base editing (BE), which can convert a single nucleotide to another, represents a newly developed and highly versatile genome engineering technology ^31; 33^. BE has advantages over other genome engineering approaches with respect to safety and clinical applicability. BE employing nickase Cas9 does not intentionally create double-stranded DNA breaks (DSBs) ^31; 33^, minimizing potential adverse effects. Also, BE with low off-targeting is being actively developed, adding an additional layer of safety ^76^. The majority of well-characterized disease-causing mutations are point mutations, and therefore many genetic disorders can be addressed by BE strategies ^77^. The robustness of BE has been demonstrated in models of genetic disorders caused by point mutations ^57; 58; 77; 78^, and the first human trial employing base editing has already been started ^79; 80^. However, many human disorders are caused by other types of mutations, such as expansions of DNA repeats ^7; 16; 81^ for which BE may not seem like an ideal tool. In contrast to this commonly held notion, we show that BE strategies could also address diseases that are caused by expanded repeats, broadening their target space and applicability.

In HD, multiple studies have shown that the uninterrupted CAG repeat length in *HTT* gene, not the polyglutamine length in huntingtin protein, determines age-at-onset ^24; 27; 30^. Age-at-onset of HD subjects carrying DI alleles not only supports this notion directly, but also points to novel therapeutic strategies. For example, converting CAG to CAA would decrease the length of uninterrupted CAG repeat without changing the length of polyglutamine or altering huntingtin protein. Indeed, our candidate BE strategies could shorten the length of uninterrupted CAG repeat by converting CAG to CAA at various sites in the CAG repeat without causing significant indels or off-target effects. In support, our candidate BE strategy modestly but significantly reduced the levels of CAG repeat expansion in mice, and HD knock-in mice carrying the CAA-interrupted repeats showed virtually zero repeat expansion. Given the role of the uninterrupted CAG repeat length as the most important disease determinant and a pivotal role for repeat instability in the modification of HD ^26; 27^, our data support the therapeutic potential of CAG-to-CAA conversion BE strategies in HD.

Our data are relevant for a number of reasons. Firstly, genetically supported targets significantly increase the success rate in clinical development ^82^; our BE strategies derive directly from human genetic observations in HD individuals ^24; 27; 30^ that point to the uninterrupted CAG repeat length in *HTT* as the most direct therapeutic target in HD. Based on these human data, CAG-to-CAA conversion even near the 5’-end of the expanded CAG repeat may produce robust onset-delaying effects. In addition, if BE strategies are applied to fully penetrant 40 or 41 CAG canonical repeats, the repeats can become reduced penetrant (i.e., 36-39). Similarly, BE strategies may be able to convert some of the reduced penetrant CAG repeats (e.g., 36 and 37 CAG) to non-pathogenic (CAG < 36), which can prevent the manifestation of the disease. Secondly, lessons from the recent huntingtin-lowering clinical trial ^83; 84^ implied the importance of allele-specific approaches ^35; 36^. Although CAG-to-CAA conversions in our experiments occurred on both mutant and normal *HTT*, our candidate BE strategies are expected to produce mutant allele*-*specific consequences. The amino acid sequence and levels of huntingtin were not altered, while the length of the uninterrupted CAG repeat was shortened. On the mutant allele, this shortening is expected to reduce the somatic instability of the repeat, reducing its disease-producing potential. On the normal allele, inherited variation in the length of the CAG repeat has not been associated with an abnormal phenotype ^19^, so the shortening of the CAG repeat is expected to be benign. Importantly, the mutant allele-specific consequences can be achieved without relying on individual genetic variations beyond the CAG repeat. Various SNP-targeting allele-specific approaches have been proposed ^35; 36; 85; 86^, but most of these can be applied only to a subset of the HD population depending on heterozygosity at the target site. Our BE strategies can achieve allele-specific consequences without targeting SNPs, and, therefore, can be applied to all HD subjects, representing a huge advantage over SNP-targeting allele-specific strategies. Lastly, BE with relaxed PAM requirements ^51; 77^ has increased the applicability of BE, and our study has further expanded the target space of this powerful technology. BE is appropriately viewed as a tool to correct disease-causing point mutations or to modify gene expression by introducing early stop codons or altering splice sites ^77; 87; 88^. However, our study demonstrates that base conversion can address disease-causing repeat expansion mutations without involving DSB. The ramifications apply not only to HD but also to numerous other diseases that are caused by expansions of repeats ^4–8^, offering alternative therapeutic approaches for the repeat expansion disorders.

Although promising, hurdles must be overcome before CAG-to-CAA conversion BE strategies are applied to humans. The BE strategies that we evaluated did not robustly generate DI alleles that are found in humans, potentially due to the possibility that PAM at specific sites that are required to generate DI did not sufficiently support the activity of CBEs that we tested. Therefore, new CBEs that can efficiently generate DI alleles will greatly facilitate the development of rational treatments for HD. Also, the inability to directly target alternative toxic species such as RAN translation or exon 1A huntingtin fragment ^89–92^ may represent one of the limitations of our BE strategies for HD. Still, if the levels of those alternative toxic species are dependent of the length of uninterrupted CAG repeat, CAG-to-CAA conversion strategies may be able to ameliorate alternative toxic species-mediated HD pathogenesis. Although BE strategies can address the primary disease driver in principle, they may not produce any significant clinical benefits if they are applied too late in the disease. As we previously speculated, the timing of treatment might have negatively impacted the outcomes of the first ASO *HTT* lowering trial ^35; 36; 83; 84^. Considering the evidence for significant levels of neurodegeneration at the onset of characteristic clinical manifestations ^93^, CAG-to-CAA conversion treatments may not produce any clinical improvements if applied late. With the expectation of mutant-specific consequences, we reason that BE strategies can be applied quite early without involving deficiency-related adverse effects ^94–97^ because CAG-to-CAA conversion is predicted to neither alter the amino acid sequence nor changes the expression levels of *HTT*. Regardless, these temporal aspects and safety features have to be determined. Finally, like other gene targeting strategies, the development of effective delivery methods is critical for applying BE therapeutically. The expansion-decreasing effects of our initial AAV injection experiments, while significant, may have been limited compared to cell culture systems by inefficient delivery and difficulty in targeting the repeat sequence. For the successful application of BE strategies to human HD, efficient delivery methods will be critical.

Given the lack of effective treatments for HD and the premature terminations of highly anticipated *HTT*- lowering clinical trials such as GENERATION-HD1 ^83^ and VIBRANT-HD (https://www.hda.org.uk/media/4418/novartis-vibrant-hd-community-letter-final-pdf.pdf), aiming at the most relevant target is becoming increasingly important. Our data reveal relevant strategies for addressing the target most strongly supported by human HD genetic data, the uninterrupted CAG repeat in *HTT*, therefore offer new opportunities for blocking the disorder at its cause. Although both great promise and significant hurdles exist for the clinical application of BE strategies in HD, our data demonstrate the proof-of-concept of this technology as the basis for developing a rational treatment for HD and, potentially, for other repeat expansion disorders.

## Abbreviations

HD: Huntington’s disease; huntingtin, *HTT*
Q: glutamine
CR: canonical repeat
LI: loss of interruption
DI: duplicated interruption
BE: base editing
CBE: cytosine base editor
PAM: protospacer adjacent motif
gRNA: guide
RNA: iPSC, induced pluripotent stem cell
EV: empty vector.

## Acknowledgements

We thank Drs. Marcy E. MacDonald, James F. Gusella, and David Liu for helpful discussion. This work was supported by grants from Harvard NeuroDiscovery Center, NIH (NS105709, NS119471, NS091161, NS049206), and CHDI Foundation. B.P.K. was also supported by an MGH ECOR Howard M. Goodman Award and a CHDI Research Agreement (14962).

## Data availability

RNAseq data of control and targeted iPSC clones have been deposited in Dryad (https://doi.org/10.5061/dryad.k3j9kd5cb).

