## Supplementary material for "Base editing strategies to convert CAG to CAA diminish the disease-causing mutation in Huntington's disease": S. Figures

**S. Figure 1. The Canonical repeat, loss of interruption, and duplicated interruption**

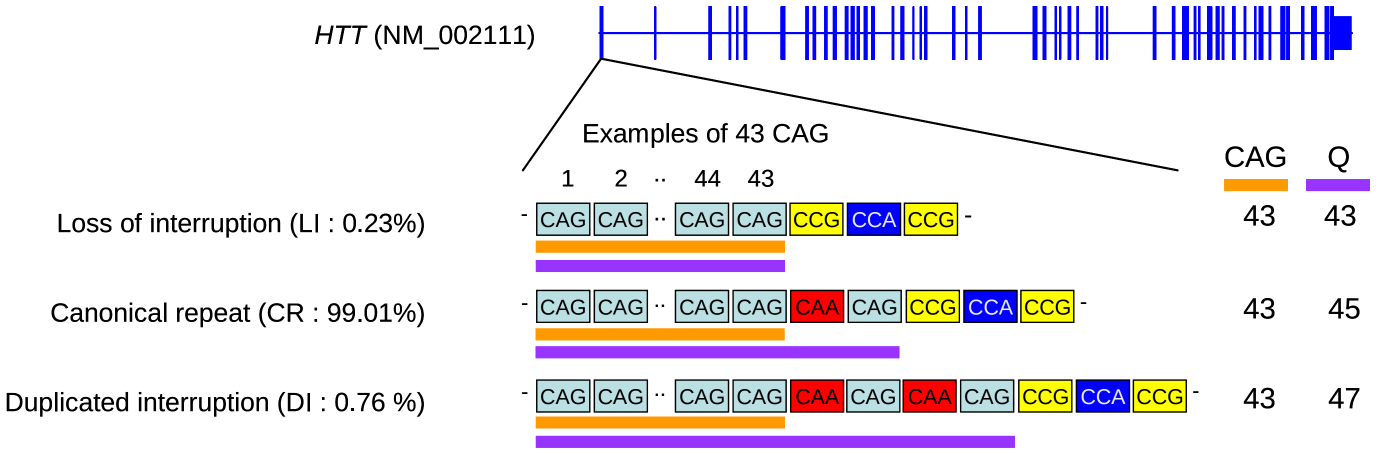


Three naturally occurring *HTT* CAG repeat alleles and their estimated frequencies are summarized. The canonical repeat (CR) consists of an uninterrupted CAG repeat followed by a CAA-CAG codon doublet; this repeat sequence is the most frequent accounting for approximately 99% of the disease chromosomes in our previous GWAS data. For CR alleles, the size of the polyglutamine stretch is longer by two than the length of the uninterrupted CAG repeat since CAA and CAG both specify glutamine (Q). Loss of interruption (LI) alleles lack the CAA interruption. This repeat is infrequent in HD subjects (0.23% in our GWAS data); the lengths of the polyglutamine stretch and uninterrupted CAG repeat are the same. Duplicated Interruption (DI) alleles, featuring two copies of the CAA-CAG, are also infrequent (0.76% in our GWA participants); the polyglutamine length is greater by four compared to the uninterrupted CAG repeat length. Orange and purple horizontal bars represent uninterrupted CAG repeat and polyglutamine, respectively. Examples were based on 43 uninterrupted CAG repeats. CAG and Q represent uninterrupted CAG repeat and polyglutamine, respectively.

**S. Figure 2. Prediction of age-at-onset of LI and DI carriers based on an assumption that polyglutamine determines onset.**

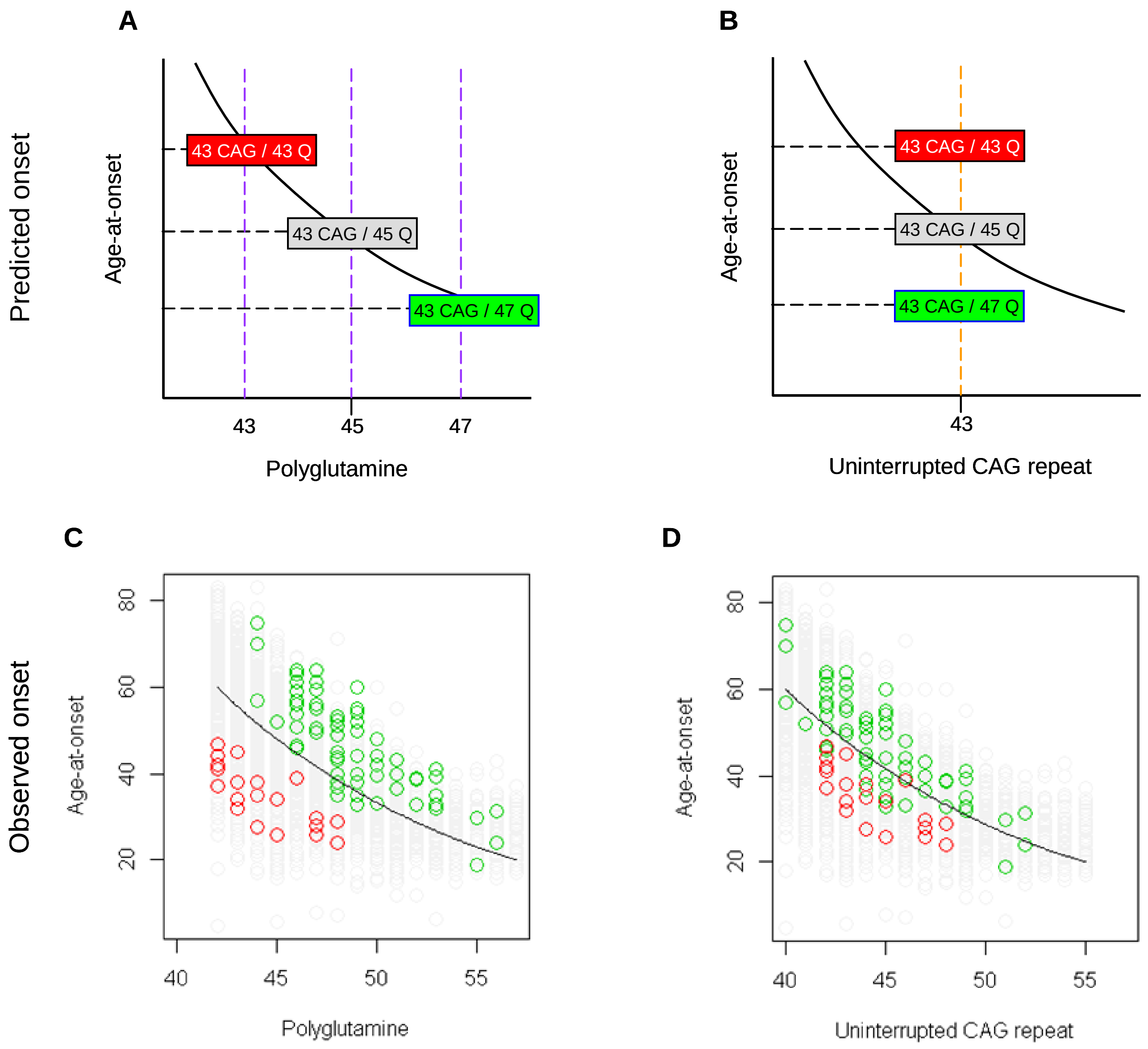


(A and B) HD subjects carrying 43 uninterrupted CAG repeats with different polyglutamine lengths due to LI and DI are displayed as examples of their positioning relative to the trend lines, which represent hypothetical regression models for CR allele carriers describing the relationship between onset age and polyglutamine length (panel A) or uninterrupted CAG repeat length (panel B). Red, grey, and green rectangles respectively represent HD subjects carrying LI, CR, and DI alleles of 43 uninterrupted CAG repeats.

(C and D) The observed age-at-onset (Y-axis) was compared to the size of polyglutamine tract (panel C) and uninterrupted CAG repeat (panel D). Green, grey, and green circles again represent HD subjects who carry DI, CR, and LI alleles, respectively. Black trend lines represent the actual regression models describing the relationship between onset age and polyglutamine length (panel C) or CAG repeat (panel D) in HD subjects carrying CR alleles. In the observed data, the LI and DI carriers showed the opposite patterns of age-at-onset from the prediction based on the assumption that polyglutamine length drives onset.

**S. Figure 3. Least squares approximation to estimate the magnitude of additional effects of LI and DI on age-at-onset.**

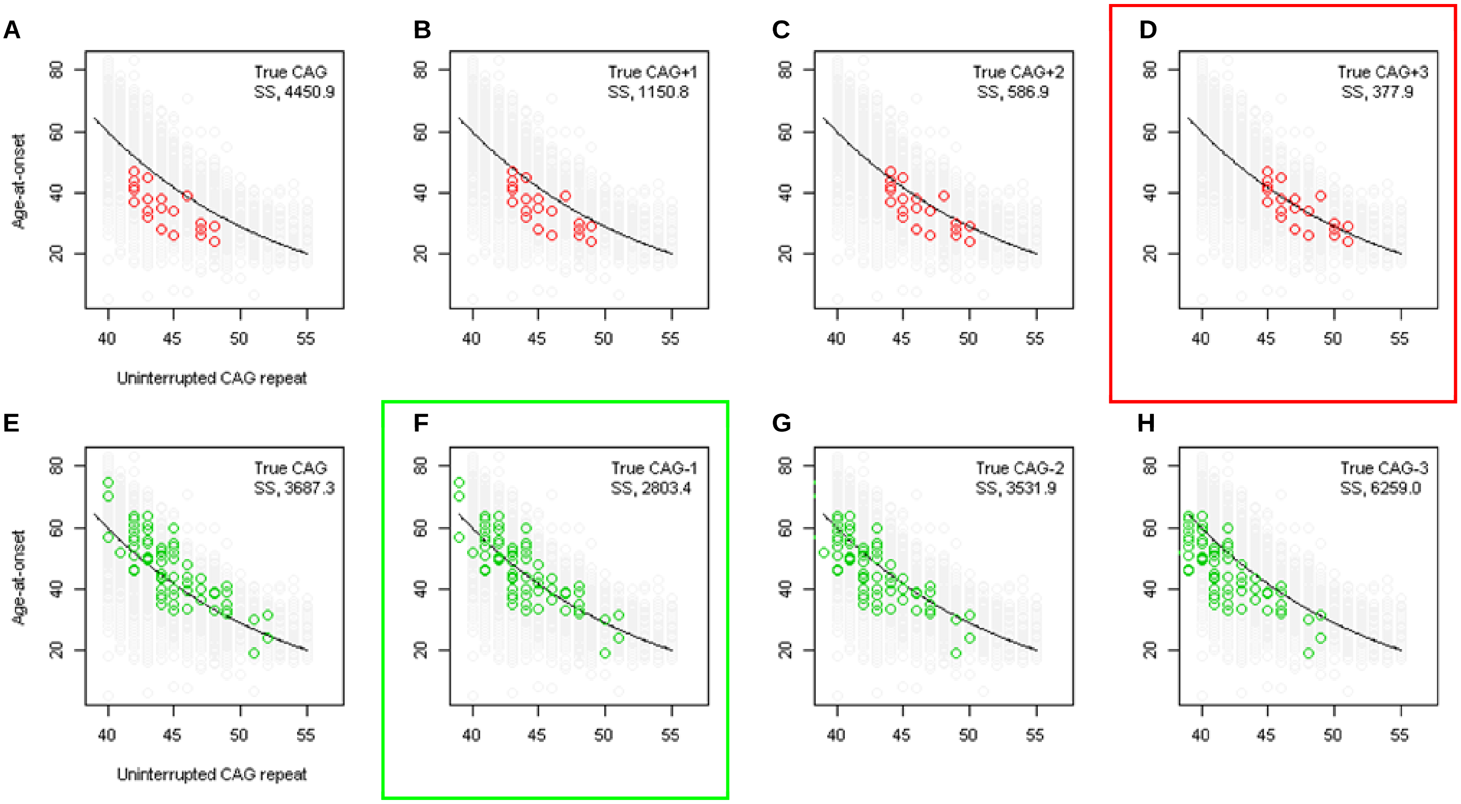


To calculate the additional effects of LI and DI alleles after accounting for uninterrupted CAG repeat length, we performed least squares approximation. For LI (red circles) carriers, we calculated the sum of square (SS) using true CAG (A), true CAG+1 (B), true CAG+2 (C) , and true CAG+3 (D). Similarly, we performed least squares approximation for DI carriers by calculating SS using true CAG (E), true CAG-1 (F), true CAG-2 (G), and true CAG-3 (H). Y-axis represents age-at-onset, and black trend lines represent the onset-CAG regression model based on CR carriers. SS values are shown at the top right corner of each plot. Least squares approximation that produced the smallest SS for LI and DI alleles are indicated by red and green rectangles, respectively.

**S. Figure 4. The gRNAs for CBEs to convert CAG to CAA in HD.**

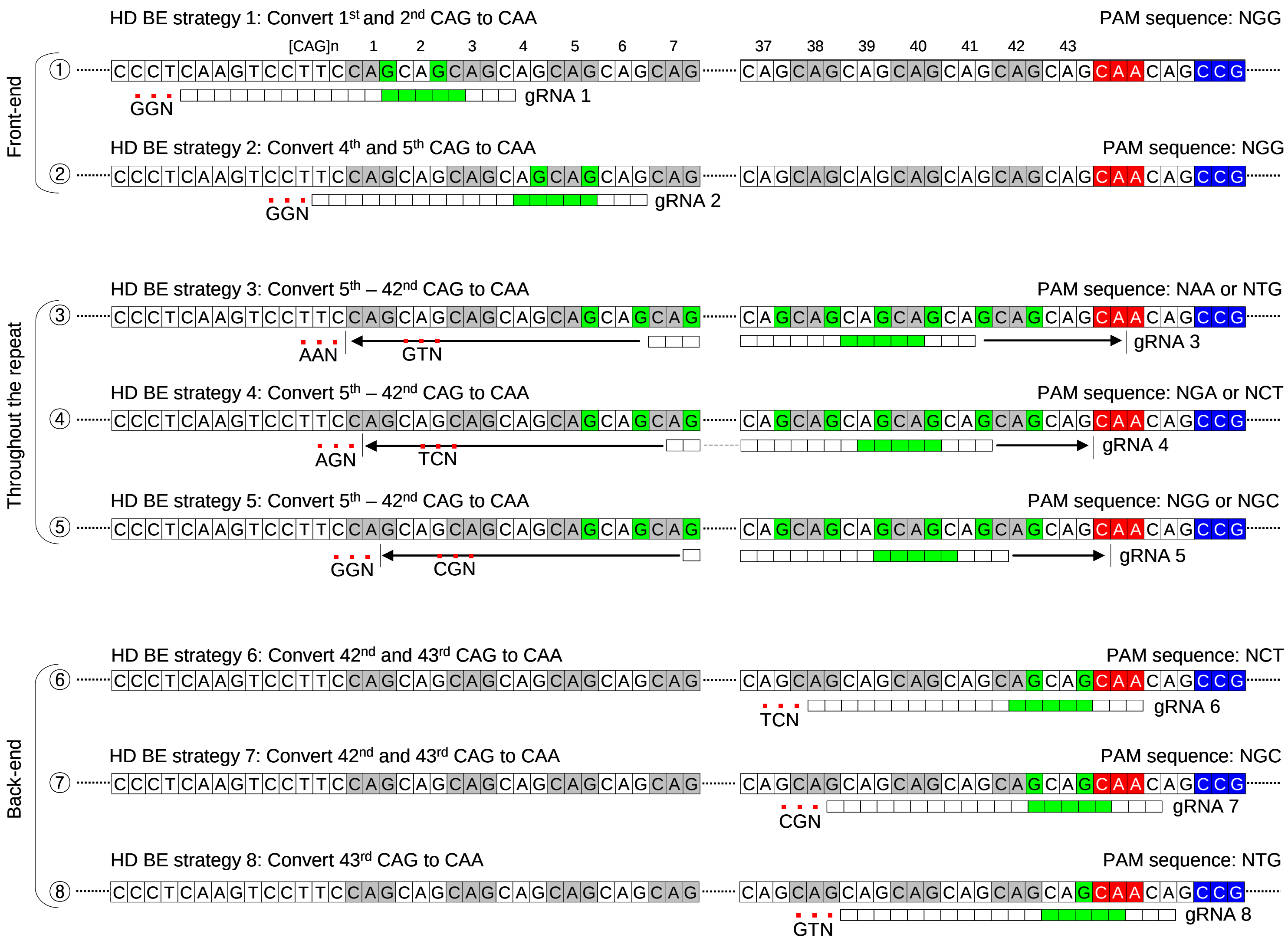


We tested eight gRNAs in this study. The gRNAs are grouped based on the predicted sites of gRNA-target hybridization. The gRNAs 1, 2 were predicted to hybridize at the front-end of the repeat; gRNAs 3, 4, and 5 were predicted to hybridize throughout the repeat; and gRNAs 6, 7, and 8 were predicted to hybridize at the back-end of the repeat. The location of the PAM (small red-filled squares under the target sequence) and gRNAs (a stretch of white and green rectangles) are indicated relative to an example of 43 CAG CR allele. The predicted conversion sites represented by the target sequence (filled green) were based on the widely used BE4.

**S. Figure 5. The lack of significant indels by BE strategies.**

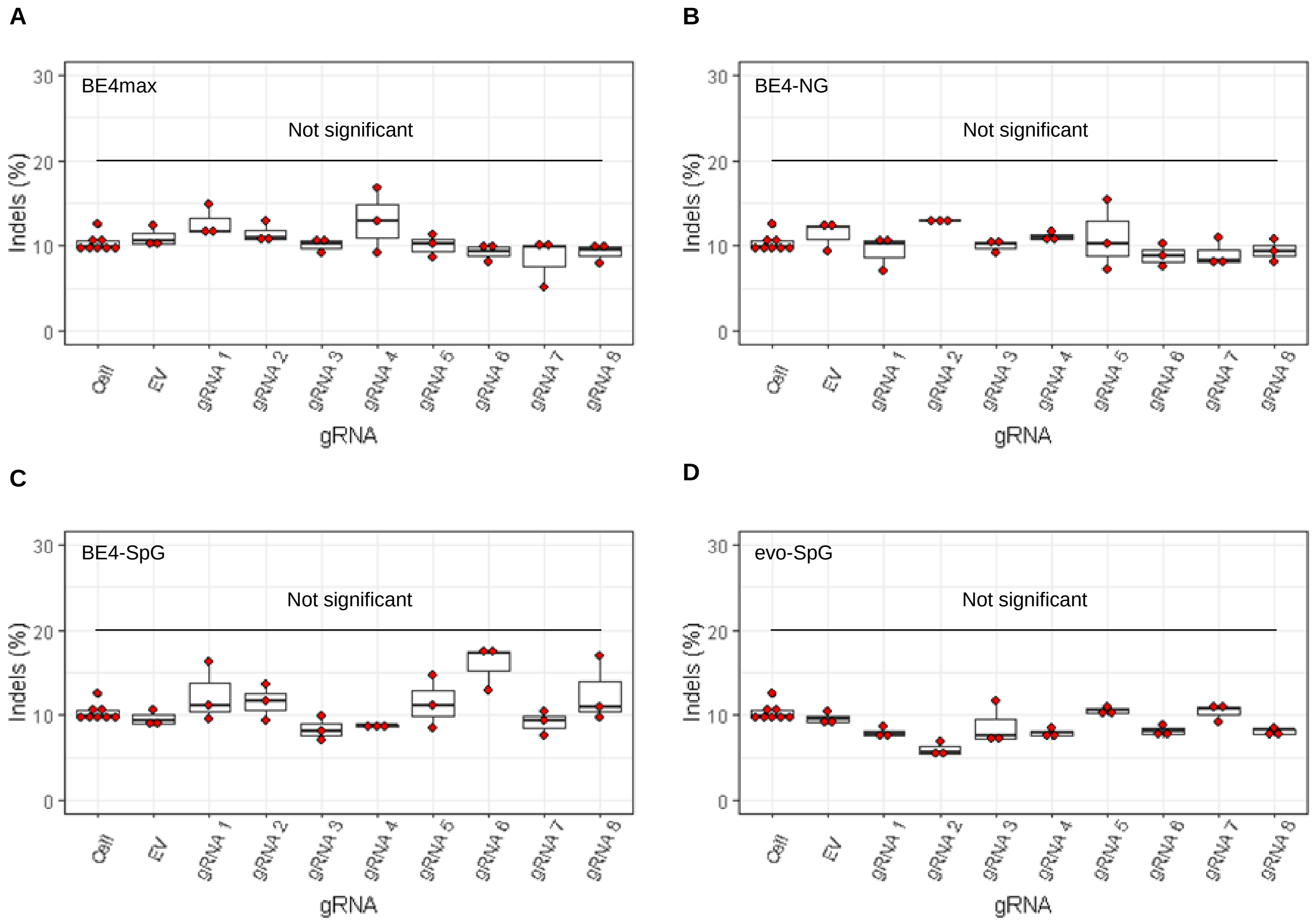


MiSeq analysis was performed on HEK293 cells treated with combinations of base editors (A, BE4max; B, BE4-NG; C, BE4-SpG; and D, evo-SpG) and gRNAs to determine the proportion of the sequence reads containing indels. Replicate samples for HEK293 cells without any treatment (i.e., Cell) were combined (n=8) and plotted for each base editor. EV represents HEK293 cells treated with a base editor and empty vector for gRNA. Each box shows the maximum, upper quarter, median, lower quarter, and minimum (from the top) based on 3 independent transfection experiments. Red circles represent individual data points. None was significant compared to corresponding EV-treated cells by Bonferroni-corrected Student t-test (8 tests for each base editor).

**S. Figure 6. Types of base conversion by CBEs.**

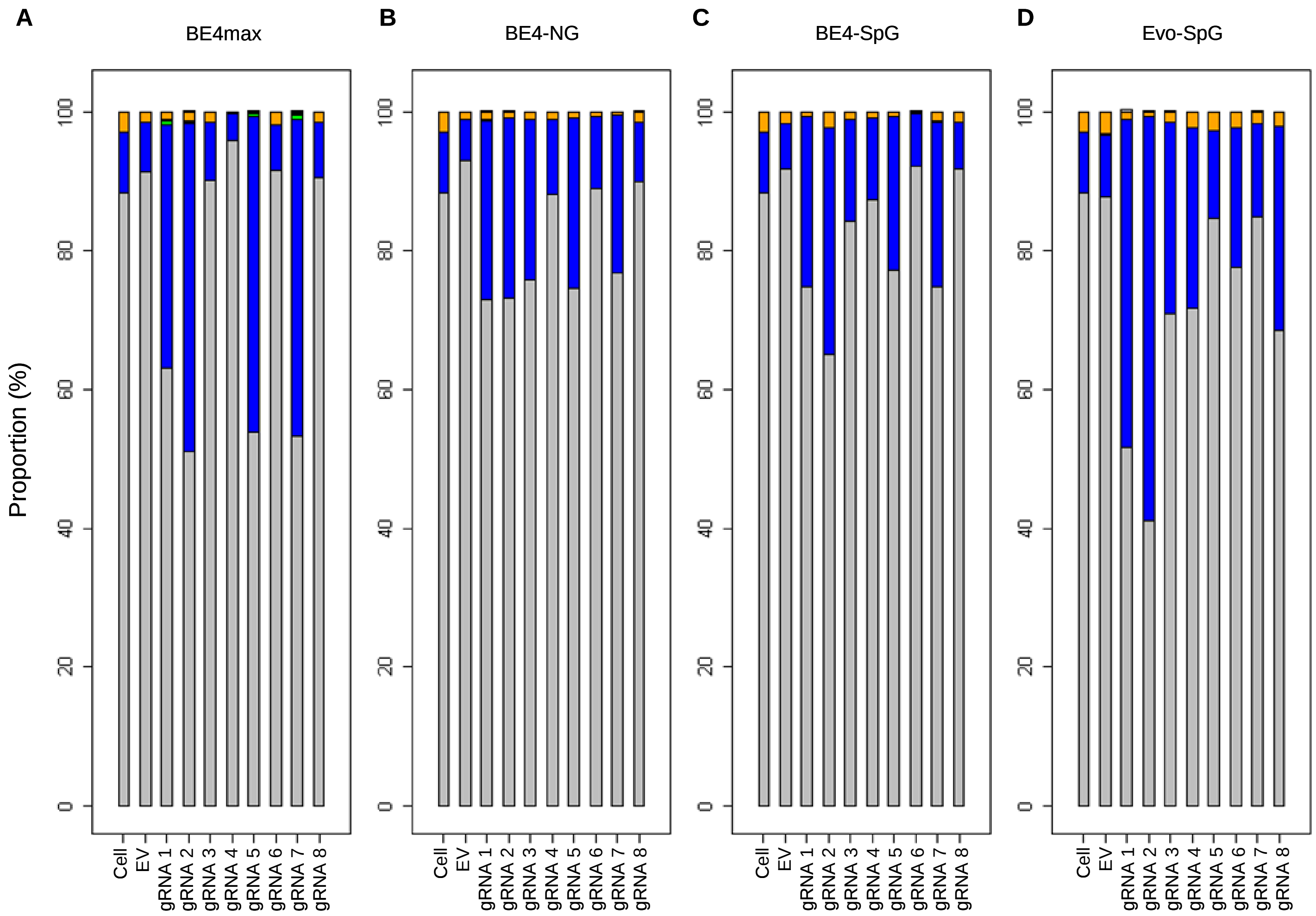


| Color | Grey | Blue | Green | Yellow | Purple | Cyan | White | Red | Black | Orange |
|---|---|---|---|---|---|---|---|---|---|---|
| Conversion | No conversion | CAA | CAC | CAT | CCG | CGG | CTG | AAG | GAG | TAG |

To determine the types of base conversions and to quantify their levels , we analyzed sequence reads of 16 or 17 CAGs (A, BE4max; B, BE4-NG; C, BE4-SpG; and D, evo-SpG). For each sample, we calculated the percentages of sequence reads containing CAA (blue), TAG (orange), and other trinucleotides. For example, the percentage of sequence reads containing CAA was calculated by dividing the number of sequence reads involving at least one CAA by the number of all sequence reads; therefore, 20% CAA means 20% of all sequence reads with 16 or 17 repeat sequences contain at least one CAA.

**S. Figure 7. Potential explanations for unexpected conversion sites.**

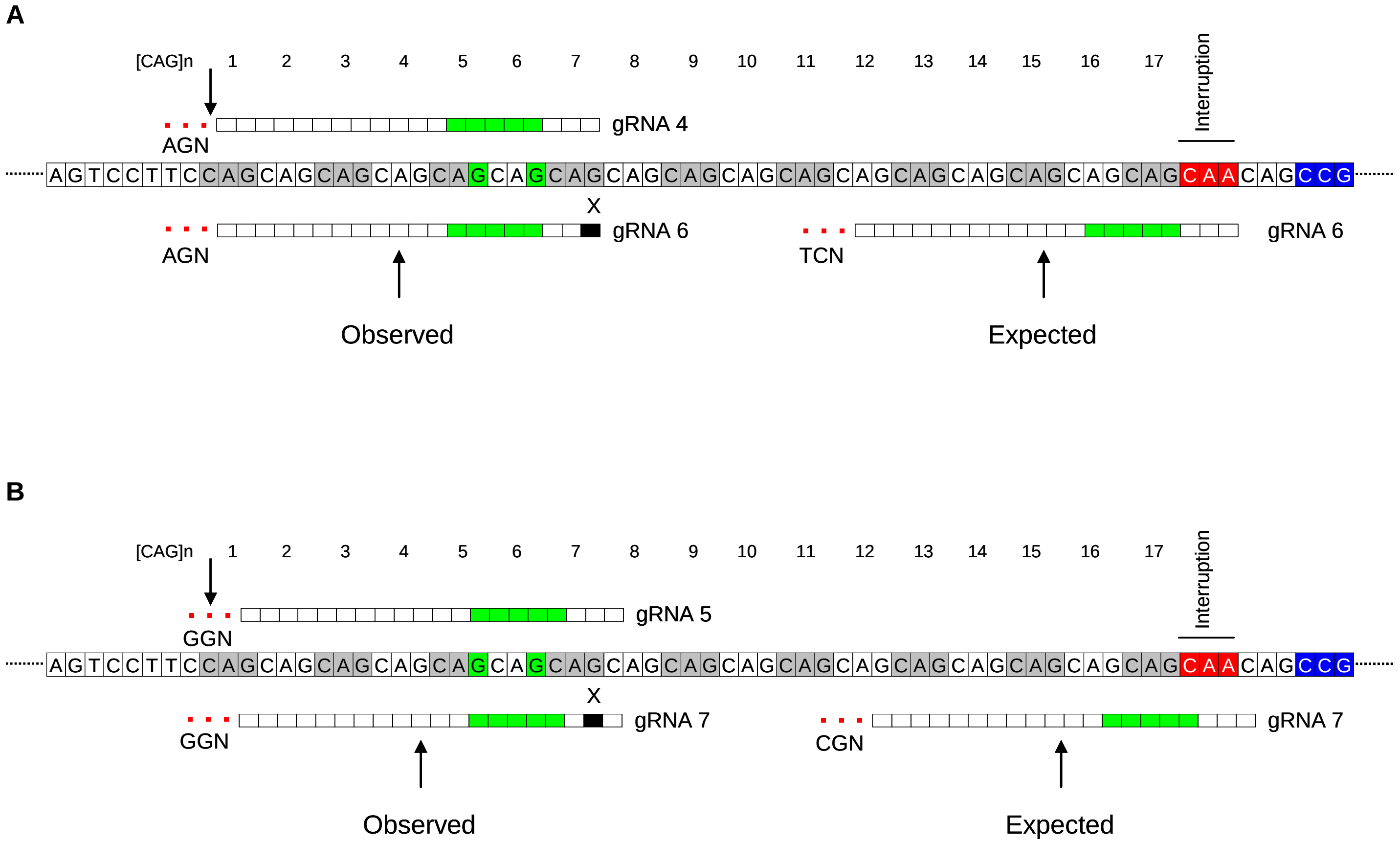


Diagrams illustrating the potential reason for the conversion at the 5th CAG by gRNA 6 (A; BE4-NG and BE4-SpG) and at the 6th CAG by gRNA 7 (b; BE4max) are shown. The sequences of gRNAs 4 and 6 are identical except for one mismatch. Therefore, it appeared that one mismatch was tolerated (black rectangle with an "X") in favor of the NGA PAM, resulting in conversion at the front end (B). Similarly, the sequences of gRNAs 5 and 7 are the same except for one nucleotide at the 19th position (B). Tolerance of one mismatch (black rectangle with an "X") would allow gRNA 7 to interact with the target site at the front-end, resulting in efficient editing at the 6th CAG, thanks to NGG PAM. Alignment and the locations of putative PAMs were indicated relative to an example of a 43 CAG canonical repeat. "G" with green highlight represents putative conversion sites; green rectangles represent conversion windows for BE4max.

**S. Figure 8. The levels of DI alleles produced by BE strategies.**

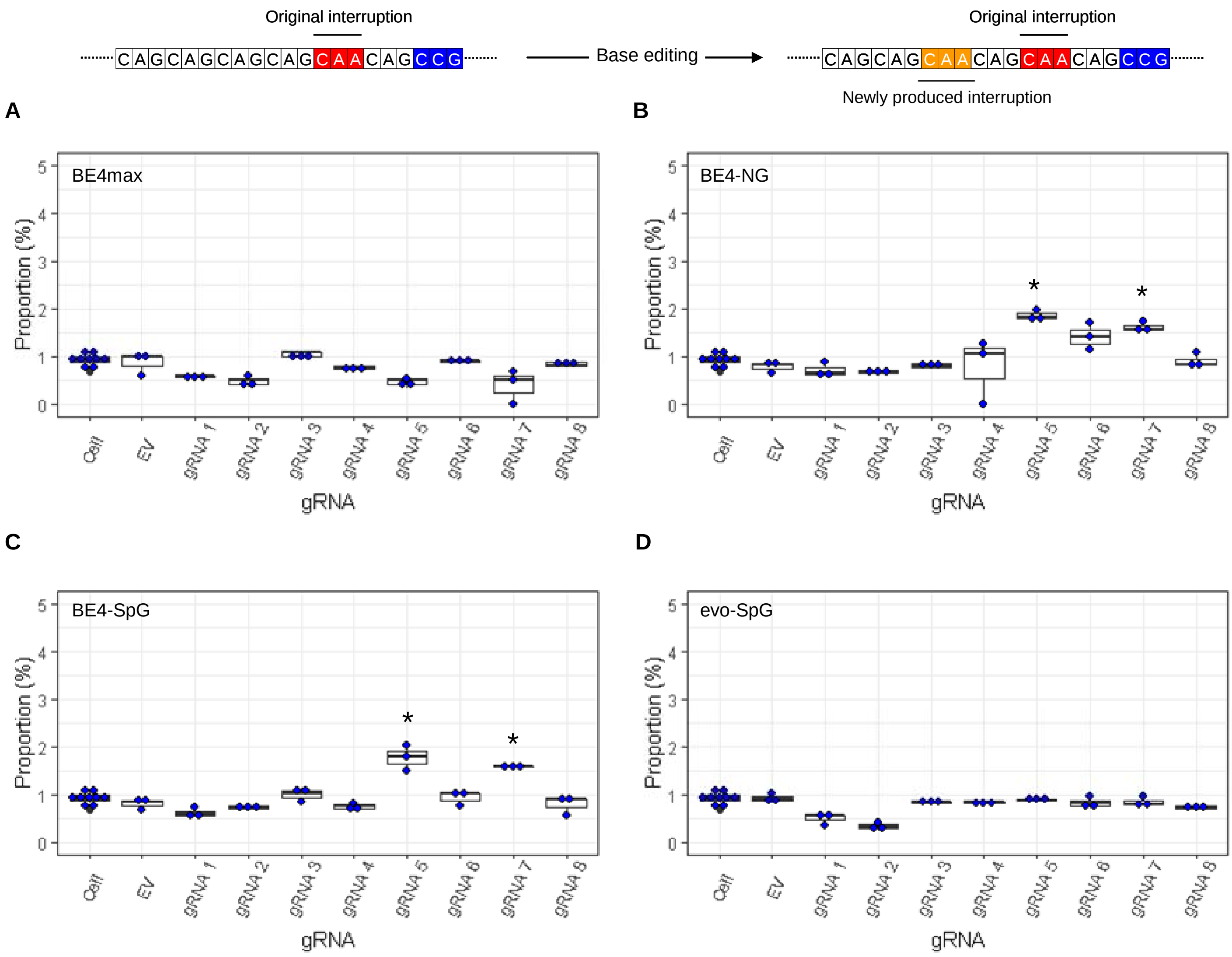


On the top, an example of duplicated interruption is displayed; filled red and orange in the example diagram represent original CAA interruption and CAA produced by BE strategies, respectively.

The proportion of DI alleles in HEK293 cells treated with different BE strategies are displayed (A, BE4max; B, BE4-NG; C, BE4-SpG; and D, evo-SpG) (n=3 independent experiments). HEK293 cells without any treatment (i.e., Cell) were combined (n=8) and plotted for each base editor. EV represents HEK293 cells treated with a base editor and empty vector for gRNA. *, significant by Bonferroni-corrected p-value < 0.05 (8 tests for each base editor).

**S. Figure 9. The levels of sequence read containing both DI and CAG-to-CAA conversions at other sites.**

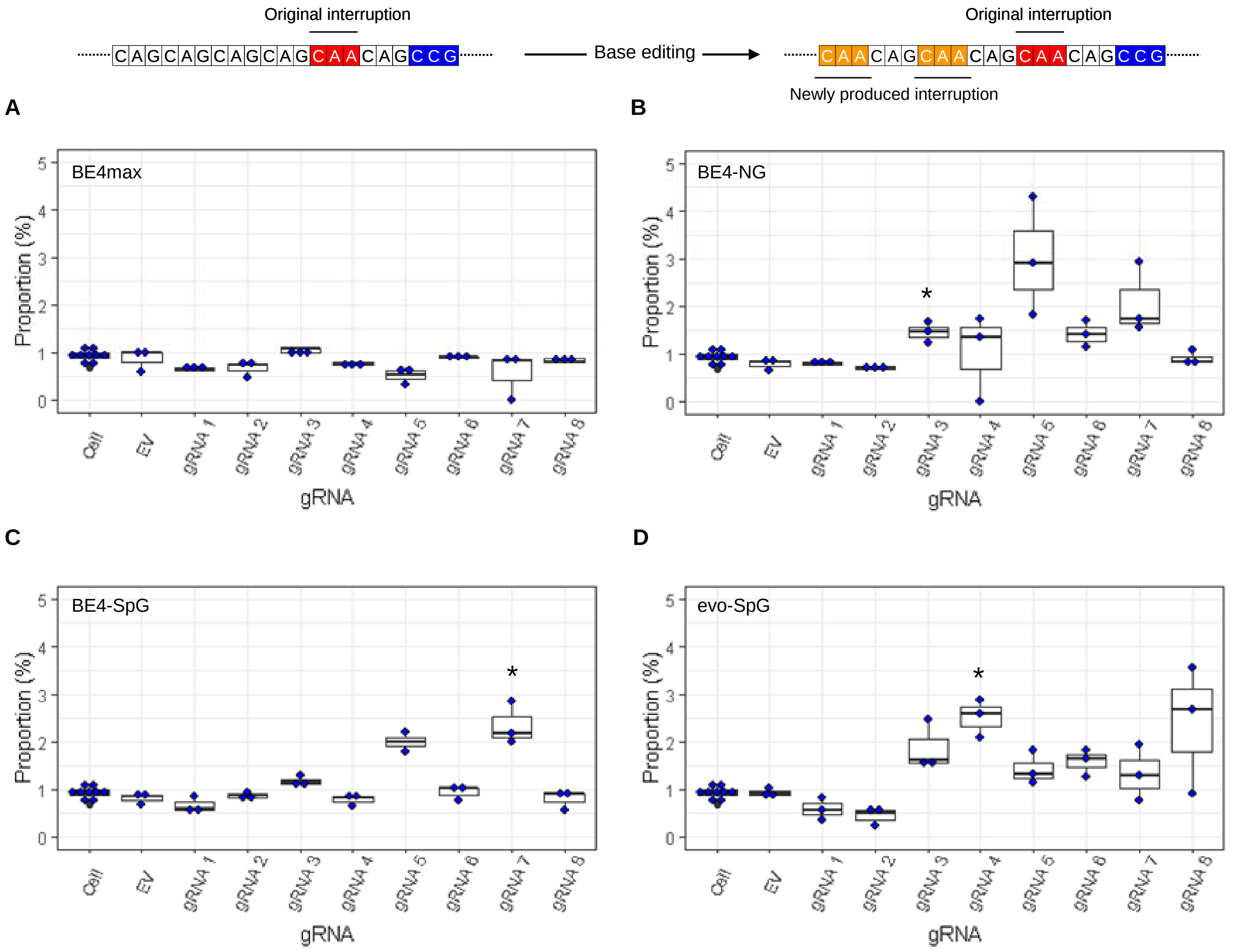


On the top, an example of DI alleles with additional CAG-to-CAA conversion is displayed; filled red and orange in the example diagram represent original CAA interruption and new CAA produced by BE strategies, respectively. The proportion of sequence reads containing DI allele and CAG-to-CAA conversions at other sites in HEK293 cells treated with different BE strategies are displayed (A, BE4max; B, BE4-NG; C, BE4-SpG; and D, evo-SpG) (n=3 independent experiments). An example of such modification is displayed at the top. HEK293 cells without any treatment (i.e., Cell) were combined (n=8) and plotted for each base editor. EV represents HEK293 cells treated with a base editor and empty vector for gRNA. *, significant by Bonferroni-corrected p-value < 0.05 (8 tests for each base editor).

**S. Figure 10. Transfection efficiency.**

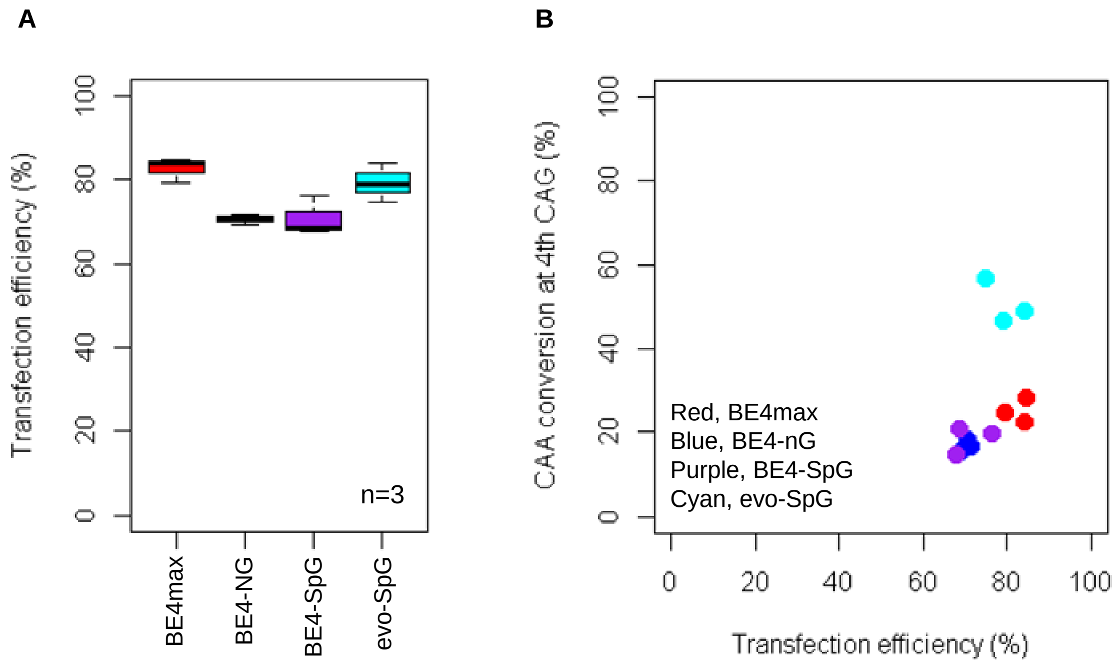


To determine whether different conversion efficiencies and patterns were due to transfection efficiencies, we treated HEK293 cells with gRNA 2, and determined transfection efficiencies (A) and conversion efficiencies (B). We compared transfection efficiency (X-axis) with the conversion efficiencies at the 4th CAG (Y-axis in the panel B). Plots were based on the mean of 3 independent transfection experiments.

**S. Figure 11. The levels of multiple conversions.**

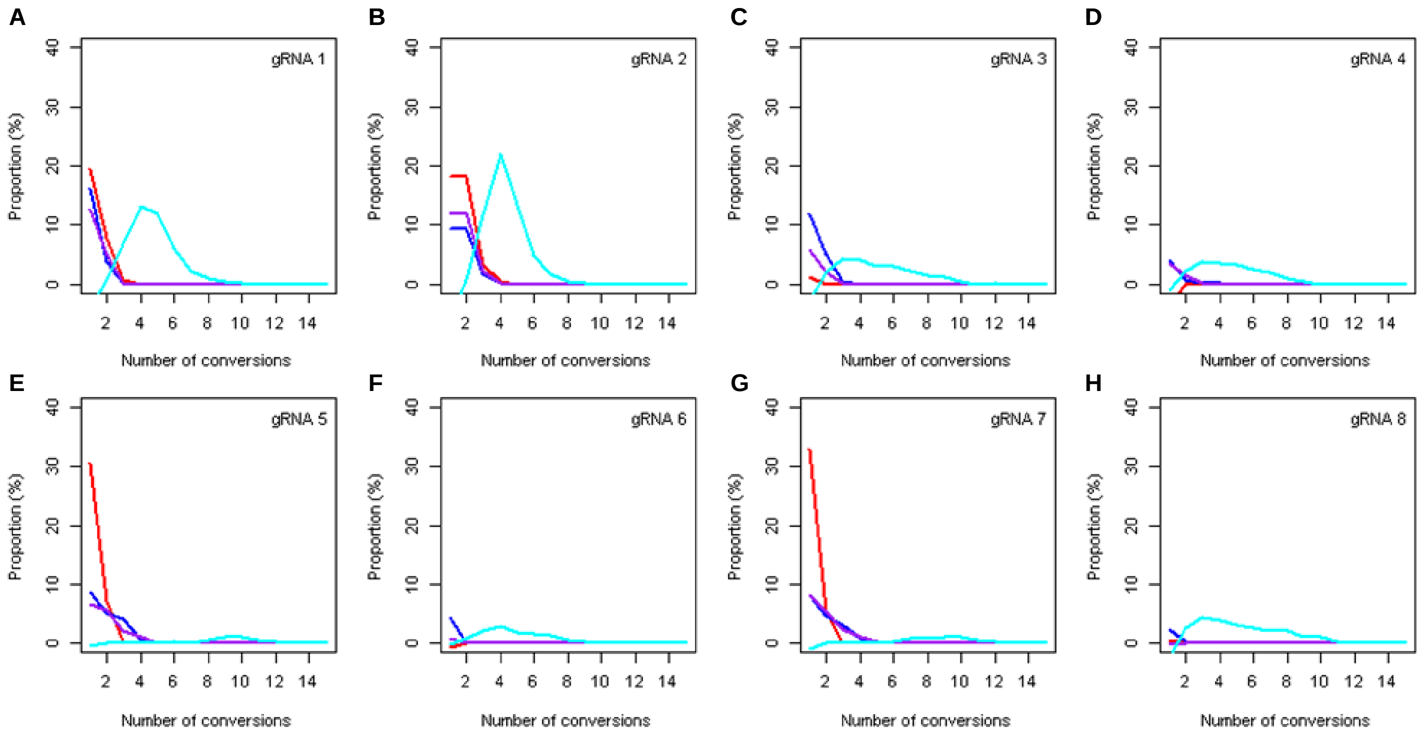


| Color | Red | Blue | Purple | Cyan |
|---|---|---|---|---|
| Base editor | BE4max | BE4-NG | BE4-SpG | evo-SpG |

The sequence reads in our MiSeq data with at least one CAG-to-CAA conversion were analyzed to count the number of CAG-to-CAA conversions in a given sequence read. The proportion was calculated by subtracting corresponding empty vector(EV)-treated cell data (Y-axis), meaning the percentage of sequence reads containing a given number of multiple conversions relative to all sequence reads from the original 16 or 17 CAG alleles. Red, blue, purple, and cyan traces represent BE4max, BE4-NG, BE4-SpG, and evo-SpG, respectively. Each panel shows a tested gRNA; plots were based on the mean of 3 independent transfection experiments in HEK293 cells.

**S. Figure 12. Off-target conversions on other polyglutamine disease genes.**

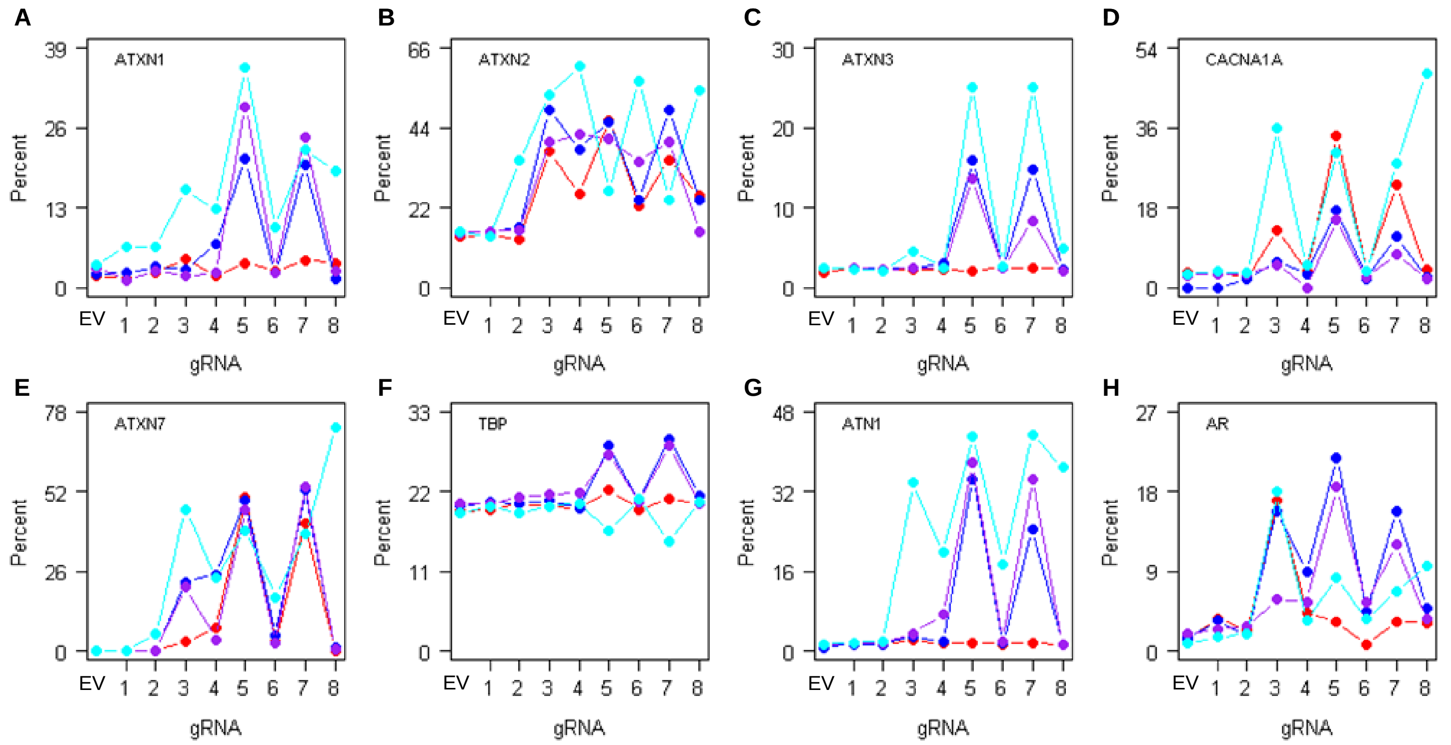


| Color | Red | Blue | Purple | Cyan |
|---|---|---|---|---|
| Base editor | BE4max | BE4-NG | BE4-SpG | evo-SpG |

To determine the levels of CAG-to-CAA conversions in the CAG repeats of other polyglutamine disease genes (A, *ATXN1*; B, *ATXN2*; C, *ATXN3*; D, *CACNA1A*; E, *ATXN7*; F, *TBP*; G, *ATN1*; and H, *AR*), we further analyzed representative HEK293 cells treated with BE strategies (n=1). EV represents empty vector-treated cells. Red, blue, purple, and cyan traces represent BE4max, BE4-NG, BE4-SpG, and evo-SpG, respectively.

**S. Figure 13. Characterization of differentiated neurons from a patient-derived iPSC.**

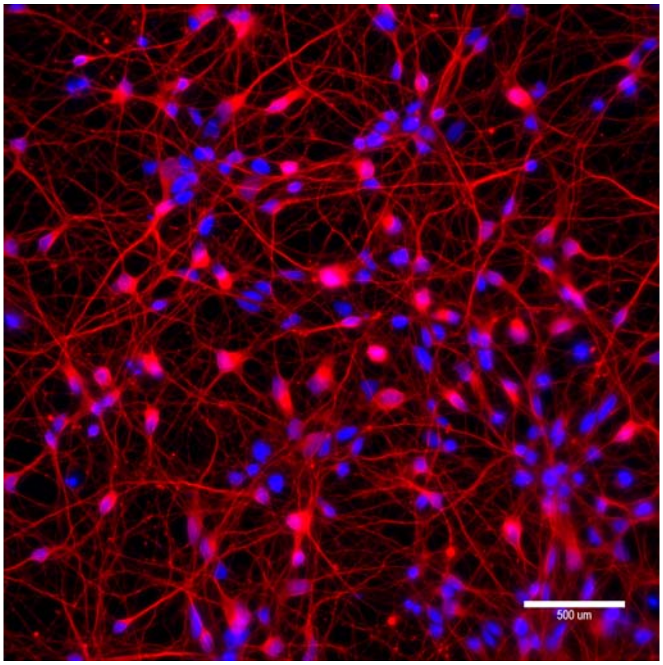


To determine the levels of allele specificities of BE strategies in neurons, we differentiated a patient-derived iPSC (42 CAG) into neurons and transfected with candidate BE strategies. Characterization of differentiated neurons was based on immunostaining of beta III tubulin (red). Blue staining represents DAPI staining for nuclei.

**S. Figure 14. Validation of HEK293-51 CAG cells.**

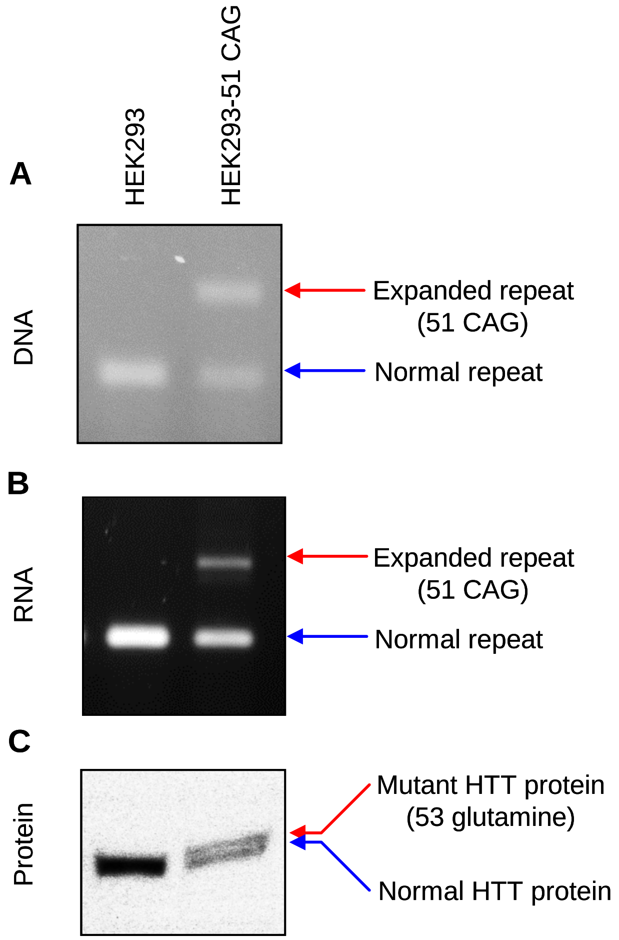


The HEK293-51 CAG cell line was generated by knocking-in a 51 CAG canonical repeat using CRISPR-Cas9. To validate the correct replacement of repeats, we analyzed DNA, RNA, and protein samples.

(A) *HTT* CAG repeat region was amplified to detect the presence of the expanded CAG repeat in the DNA.

(B) Similarly, RNA samples were amplified focusing on the CAG repeat region to determine whether HEK293-51 CAG cells express *HTT* mRNA harboring an expanded repeat.

(C) Immunoblot analysis was performed to confirm the expression of mutant HTT protein. Note, the separation between mutant and normal HTT protein was marginal because the size difference between the two is relatively very small (C). HEK293 and HEK293-51 CAG represent original carrying 16/17 CAG canonical repeats and HEK293 cells carrying 17/51 CAG canonical repeats, respectively.

**S. Figure 15. The lack of significant alterations in gene expression by BE4max-gRNA 1 and BE4max-gRNA 2.**

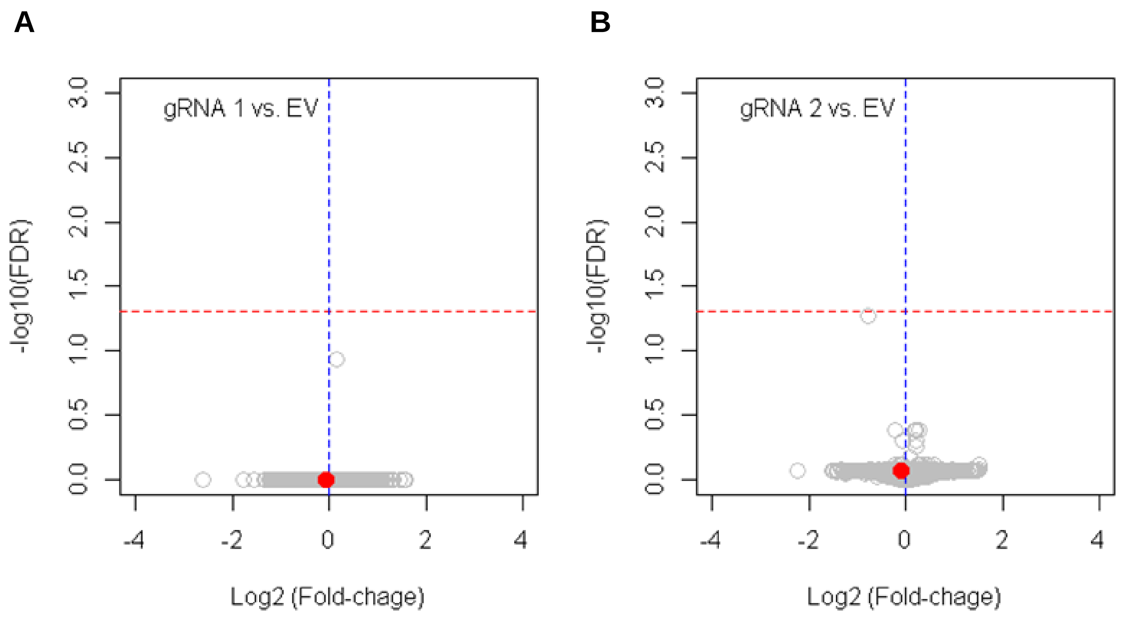


HEK293 cells were treated with empty vector (EV), or candidate BE strategies such as BE4max-gRNA 1 (A), and BE4max-gRNA 2 (B). Subsequently, DNA samples and RNA samples were collected for MiSeq analysis and RNAseq analysis to evaluate the levels of on-target conversion and changes in transcriptome (B and C; n=4), respectively. The most significant gene in cells treated with BE4max-gRNA 2 (panel B) was *HSD3B1*, which was not significant by false discovery rate of 0.05 (red lines).
